# The transcription factor Zic4 acts as a transdifferentiation switch

**DOI:** 10.1101/2021.12.22.473838

**Authors:** Matthias Christian Vogg, Jaroslav Ferenc, Wanda Christa Buzgariu, Chrystelle Perruchoud, Panagiotis Papasaikas, Paul Gerald Layague Sanchez, Clara Nuninger, Céline Delucinge-Vivier, Christine Rampon, Leonardo Beccari, Sophie Vriz, Stéphane Vincent, Brigitte Galliot, Charisios D. Tsiairis

## Abstract

The molecular mechanisms that maintain cell identities and prevent transdifferentiation remain mysterious. Interestingly, both dedifferentiation and transdifferentiation are transiently reshuffled during regeneration. Therefore, organisms that regenerate readily offer a fruitful paradigm to investigate the regulation of cell fate stability. Here, we used *Hydra* as a model system and show that *Zic4* silencing is sufficient to induce transdifferentiation of tentacle into foot cells. We identified a Wnt-controlled Gene Regulatory Network that controls a transcriptional switch of cell identity. Furthermore, we show that this switch also controls the re-entry into the cell cycle. Our data indicate that maintenance of cell fate by a Wnt-controlled GRN is a key mechanism during both homeostasis and regeneration.

**One-Sentence Summary:** A Wnt-controlled GRN controls fate maintenance in *Hydra*.

## Main Text

When cells differentiate during embryonic development or adult regeneration, they acquire specific characteristics, a specialization compensated by a loss of plasticity (Lambert et al. 2021). The protection of the identity of terminally differentiated cells relies on mechanisms that are poorly understood. Therefore, model systems that readily reactivate developmental processes such as the freshwater polyp *Hydra*, help decipher the mechanisms underlying the balance between plastic cell fates and stable cell identities. The animals, which can regenerate any missing body part after amputation, are composed of two epithelial cell layers, the epidermis and the gastrodermis, which in the body column are populated with three distinct stem cell populations, the epidermal and gastrodermal epithelial stem cells (ESCs), and the multipotent interstitial stem cells (ISCs) (Vogg et al. 2021). The ESC populations proliferate continuously in the body column, and terminally differentiate when they reach the apical (head) or basal (foot) regions. The tip of the head, named hypostome, harbors a head organizer required to maintain apical patterning in intact animals, an activity positively regulated by Wnt3/β-catenin signaling and restricted by the transcription factor Sp5 (Nakamura et al. 2011)(Vogg et al. 2019) (**Figure 1A**). When *Sp5* expression is knocked-down, *Wnt3* is de-repressed along the body column and ectopic head formation occurs. The base of the head is surrounded by tentacles where the epidermal layer is made of large epithelial cells arrested in G2 and terminally differentiated as tentacle battery cells (TBC) (**Figure Supplement 1**). Each TBC contains approximately twenty mechano-sensory cells named nematocytes (or stinging cells). These cells discharge a venom capsule named nematocyst when a specialized cilium, the cnidocil, is stimulated by prey (Holstein 2012). At the other extremity of the animal, the epidermal epithelial cells of the basal disc (foot) named basal disc cells (BDC) are terminally differentiated and produce acid mucopolysaccharide (MPS) droplets that facilitate the attachment to substrates (Rodrigues et al. 2016). Thus, both the tentacles and the foot display highly differentiated cells. Whether terminally differentiated epithelial cells can acquire an alternative identity in *Hydra* is unknown. Here, we show that the silencing of the transcription factor *Zic4* is sufficient to induce transdifferentiation of tentacle battery cells into basal disc cells.

**Figure 1.**
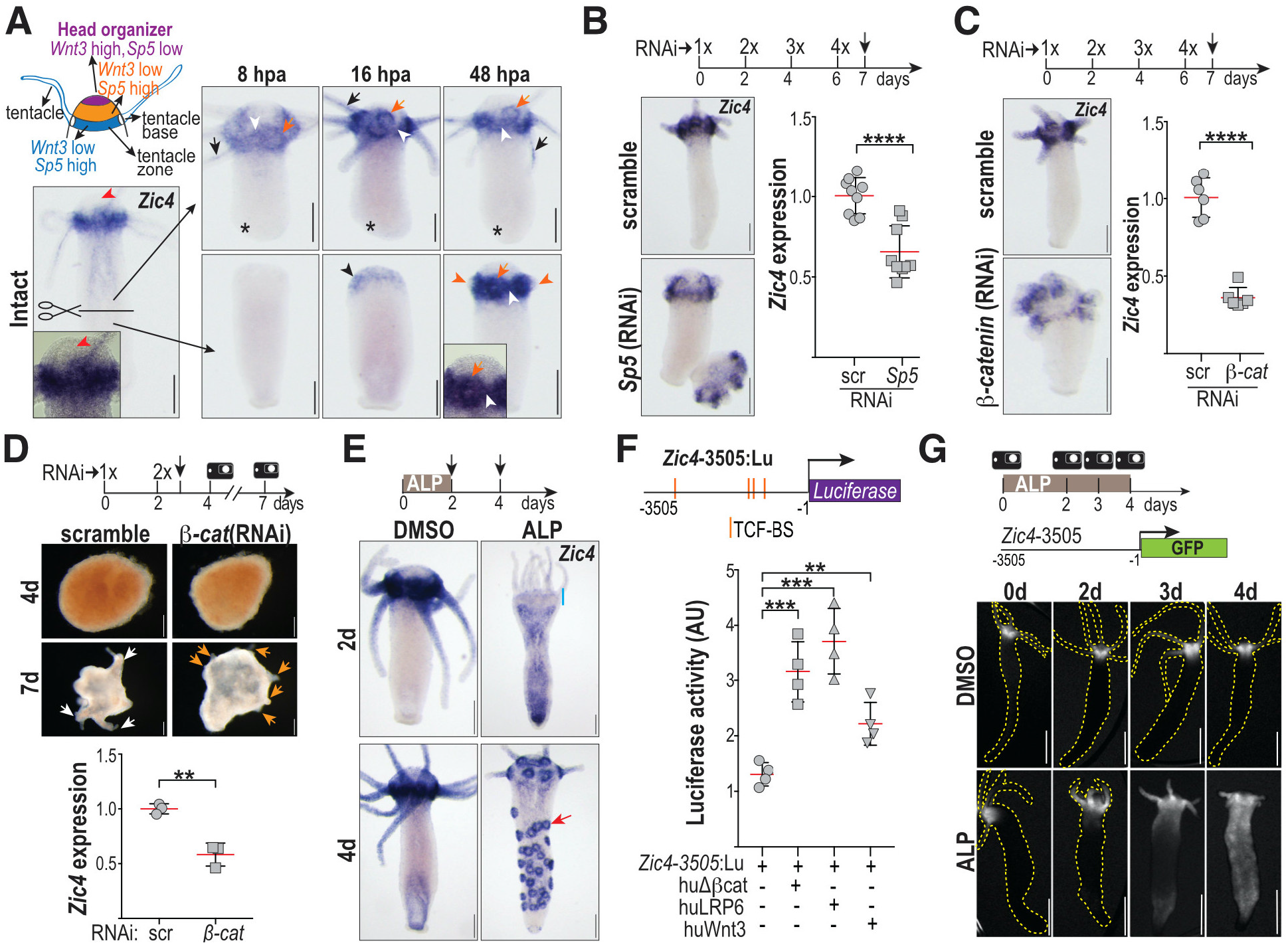
*Zic4*, a tentacle zone marker, target of Sp5 and Wnt/β-catenin signaling. **(A)** Scheme depicting *Wnt3* and *Sp5* apical expression domains. *Zic4* expression detected in homeostatic heads: tentacles (black arrows), tentacle base (orange arrows), tentacle zone (white arrowheads), hypostome (red arrowheads), during basal (asterisks) and apical regeneration at 16 hpa (black arrowhead) and 48 hpa (orange arrowheads: tentacle rudiments). **(B-C)** *Zic4* expression detected by ISH and qPCR (d7) after 4 electroporations with scramble (control), *Sp5* or *β-catenin* siRNAs. **(D)** Apical differentiation in reaggregates obtained from animals electroporated twice with scramble or *β-catenin* siRNAs before dissociation (black arrow): apical rudiments (orange arrows), fully differentiated heads (white arrows). (Bottom) *Zic4* expression measured by qPCR in reaggregates taken at day-7. **(E)** Ectopic *Zic4* expression in ALP-treated animals, reduced at the apex (blue line) at day-2, in tentacle bases (red arrow) at day-4. **(F)** Luciferase activity driven by the *Zic4* promoter in HEK293T cells when β-catenin signaling is activated. Orange bars: TCF binding-sites (TCF-BS). **(G)** GFP fluorescence in *Zic4-3505:GFP* transgenic animals exposed to ALP. In B, C, D and F, each point represents one biological replicate. Statistical p-values: **≤ 0.01; ***≤0.001; ****≤ 0.0001. Error bars indicate SD. Scale bars: 200 μm.

Among 83 putative *Sp5* target genes previously identified in HEK293T cells (Vogg et al. 2019), we focused in the zinc-finger transcription factor *Zic4* that displays an apical-to-basal graded expression along the *Hydra* body axis. Interestingly, *Zic4* is predominantly expressed in the tentacle zone of intact animals and re-expressed within 16 hours after amputation in apical-regenerating tissues (**Figure 1A, Figure Supplement 2-3, Dataset Supplement 1**). Amongst the four *Hydra Zic*-related genes, *Zic4* is the only one to show these features (**Figure Supplement 4-5**). *Zic4* appears positively regulated by *Sp5* and Wnt/β-catenin signaling as its expression is notably reduced in the tentacle zone after *Sp5* or *β-catenin* silencing (**Figure 1B-1D, Figure Supplement 6**). When Wnt/β-catenin signaling is ubiquitously activated by Alsterpaullone (ALP) treatment, a dual regulation is observed for *Zic4*, a down-regulation in the apical region together with a global up-regulation in the body-column after two days. This turns into an “Octopus” phenotype after four days, i.e. multiple *Zic4*-expressing rings, with each ring corresponding to the base of an ectopic tentacle (**Figure 1E, Figure Supplement 7**). This regulation is likely direct as the 3’505 bp *Zic4* upstream sequence contains four consensus TCF/LEF binding sites (**Figure Supplement 8**). When expressed in HEK293T cells, the *Zic4-3505*:luciferase construct is up-regulated when human β-catenin, Wnt3 or LRP6 are overexpressed (**Figure 1F**). In zebrafish embryos the overexpression of *HyZic4* induces Wnt-like phenotypes, namely eye and axial anomalies, which reflect inappropriate Wnt/β-catenin signaling (**Figure Supplement 9**). To assess *Zic4* regulation *in vivo*, we produced a *Zic4-3505:GFP* transgenic line, and recorded an increase in *GFP* expression following ALP treatment (**Figure 1G**). These data demonstrate a positive regulation of Wnt/β-catenin signaling and Sp5 on *Zic4* expression in the tentacle zone in homeostatic conditions, and along the body column when Wnt/β-catenin signaling is up-regulated.

To explore Zic4 function, we knocked-down *Zic4* by RNA interference (RNAi). Intact *Zic4*(RNAi) animals exhibit tentacles with half the length of those in control animals. The overall tentacle number is not affected (100%; *n* = 100) (**Figure 2A, Figure Supplement 10A**). In contrast, apical-regenerating *Zic4*(RNAi) animals, regenerate shorter but also 25% fewer tentacles (100%; *n* = 99) **(Figure 2B, Figure Supplement 10B**). We also knocked-down *Zic4* in ALP-treated *Hydra* and found the development of ectopic tentacles along the body column strongly impaired (**Figure 2C, Figure Supplement 10C**). Consistent with a role in tentacle formation, we also observed ectopic *Zic4* expression in the second tentacle ring induced by DAC-2-25 treatment (Glauber et al. 2013) (**Figure Supplement 11**). We also noticed that the expression of two components of the head organizer, *Wnt3* and *Bra1*, is not affected in intact or regenerating *Zic4*(RNAi) animals, suggesting that *Zic4* is required neither for the maintenance nor for the formation of a functional head organizer (**Figure 2D, Figure Supplement 12**). Indeed, when we transplanted apical tissue from control and *Zic4*(RNAi) animals into *actin*:GFP transgenic animals, we observed a similar induction of a secondary body axis, a hallmark of a functional head organizer (**Figure Supplement 13**). We concluded that *Zic4* is required for tentacle maintenance and tentacle formation.

**Figure 2.**
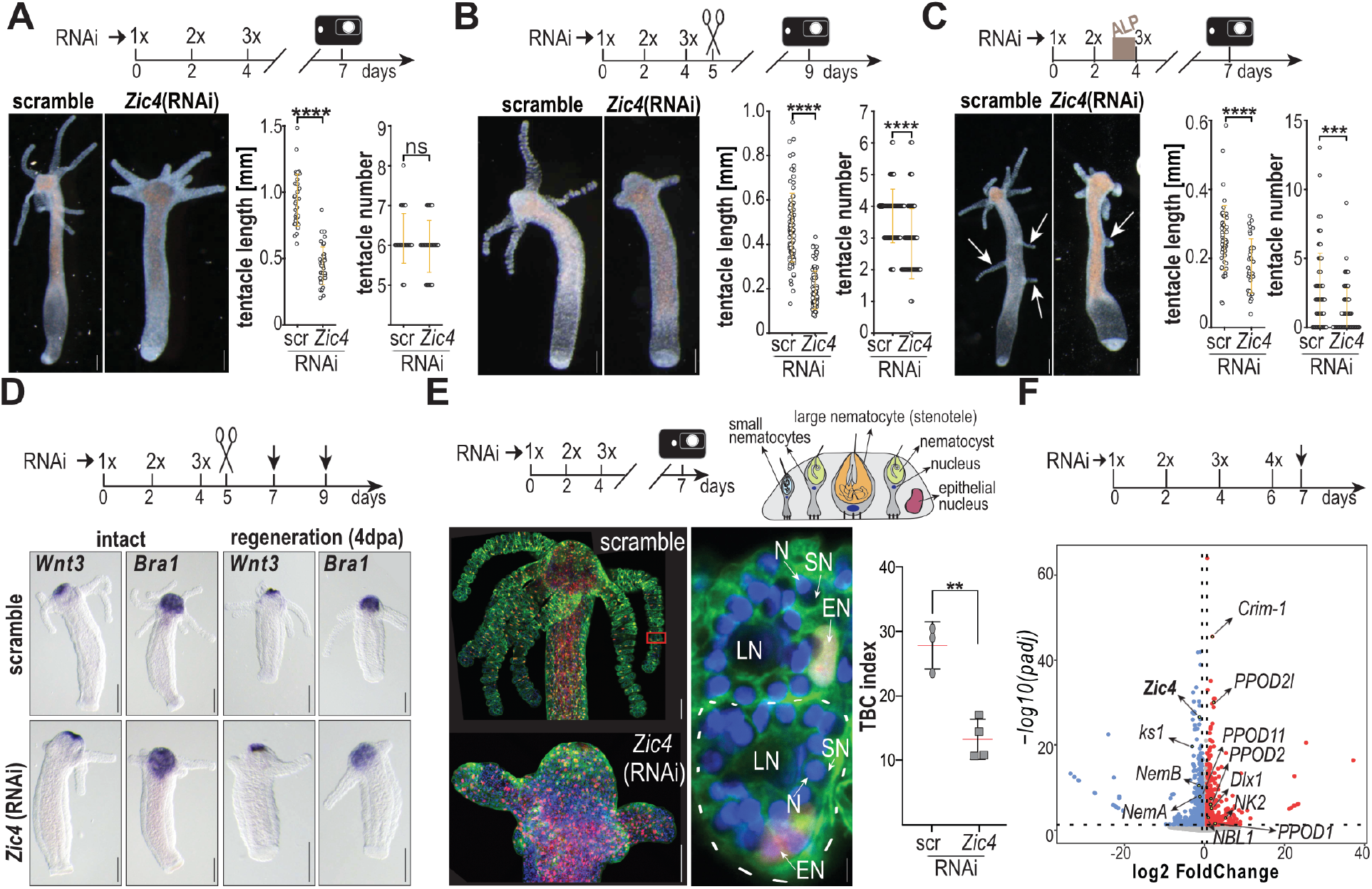
Zic4 is required for Tentacle Battery Cell (TBC) differentiation and tentacle formation. **(A-C)** Impact of *Zic4* silencing in intact (A), apical-regenerating (B) and ALP-treated (C) *Hv_Basel* animals. *Zic4*(RNAi) animals exhibit shorter (A-C) and fewer tentacles (B-C). Each data point represents one animal. White arrows: ectopic tentacles. **(D)** *Wnt3* and *Bra1* expression in intact (left) and head-regenerating (right) *Zic4*(RNAi) *Hv_Basel* fixed on d7 (intact) and d9 (apical-regenerating). **(E)** Schematic view of a tentacle battery cell (TBC) that typically contains one large nematocyte (LN, stenotele) and numerous small nematocytes (SN). Epidermal epithelial cells and TBCs from transgenic FUCCI-eGFP *Hydra* constitutively express cytoplasmic GFP and nuclear mCherry. Red square: enlarged TBCs, one being outlined with a white dashed line, EN: epithelial nuclei; N: DAPI-stained nuclei. The TBC index corresponds to the number of TBCs per 100 gastrodermal cells counted on macerated apical regions from scramble and *Zic4*(RNAi) animals. Each point represents one replicate from two independent experiments. **(F)** Volcano plot representing differentially expressed genes after *Zic4*(RNAi) as indicated. Statistical p-values: **≤ 0.01; ***≤0.001; ****≤ 0.0001. Error bars indicate SDs. Scale bars: 200 μm (A-D), 100 μm (E), 5 μm for enlarged views.

Next, we searched for changes underlying deficient tentacle maintenance and tentacle formation in *Zic4*(RNAi) animals. To monitor the cellular composition of tentacles, we used transgenic FUCCI-*Hydra* that constitutively express GFP in epidermal epithelial cells and noted a two-fold decrease in the TBC number after *Zic4*(RNAi) (**Figure 2E, Figure Supplement 14-15**). Nevertheless, we found the levels of epithelial proliferation and epithelial apoptosis unchanged after *Zic4* or *Zic4/Sp5* knock-down (**Figure Supplement 16**). Similarly, we did not record any change in the migration/displacement behavior of cells towards the head region (**Figure Supplement 17**). The RNA-seq analysis performed on scrambled and *Zic4*(RNAi) animals **(Figure 2F, Dataset Supplement 2**) reveals a striking decrease in the transcript level of three tentacle markers *ks1, nematocilin-A, nematocilin-B* (Hwang et al. 2008)(Hwang et al. 2007), consistent with the *Zic4*(RNAi)-induced loss of TBCs. Furthermore, we noted an up-regulation of genes normally expressed in the basal disc, including the peroxidases *PPOD1, PPOD2, PPOD2l, PPOD11*, the homeogenes *Dlx1* and *NK2*, the transmembrane BMP regulator *Crim-1* and the BMP antagonist *NBL1* (Wenger et al. 2019) (**Figure 2F, Table Supplement 2**). These findings suggest that *Zic4* controls tentacle identity by repressing basal gene expression.

In *Zic4*(RNAi)*, Sp5*(RNAi) and *Zic4/Sp5(RNAi)* animals, we found tentacle regions that express the basal marker *Crim-1* but no longer *Nematocilin-A*, as well as ectopic structures along the body column that do not express *Wnt3* but express *Crim-1* three days post-EP3 (**Figure 3A, Figure Supplement 18-21**). In the same conditions, we observed MPS droplets typical of basal disc cells (BDC) in tentacle cells as well as in the structures ectopically formed along the body column when *Zic4* and/or *Sp5* are knocked-down (**Figure 3B, Figure Supplement 20-21**). This phenotype is enhanced in *Zic4/Sp5(RNAi)* animals that develop complete ectopic basal discs, which are morphologically indistinguishable from those of control animals (100%, *n* = 70). We observed a similar tentacle to basal disc transformation upon *β-catenin* silencing (**Figure Supplement 22**), likely as a result of *Zic4* down-regulation. This tissue transformation exhibits a stronger penetrance in *Hv_AEP* than in *Hv_Basel* animals (**Figure Supplement 18-23**).

**Figure 3.**
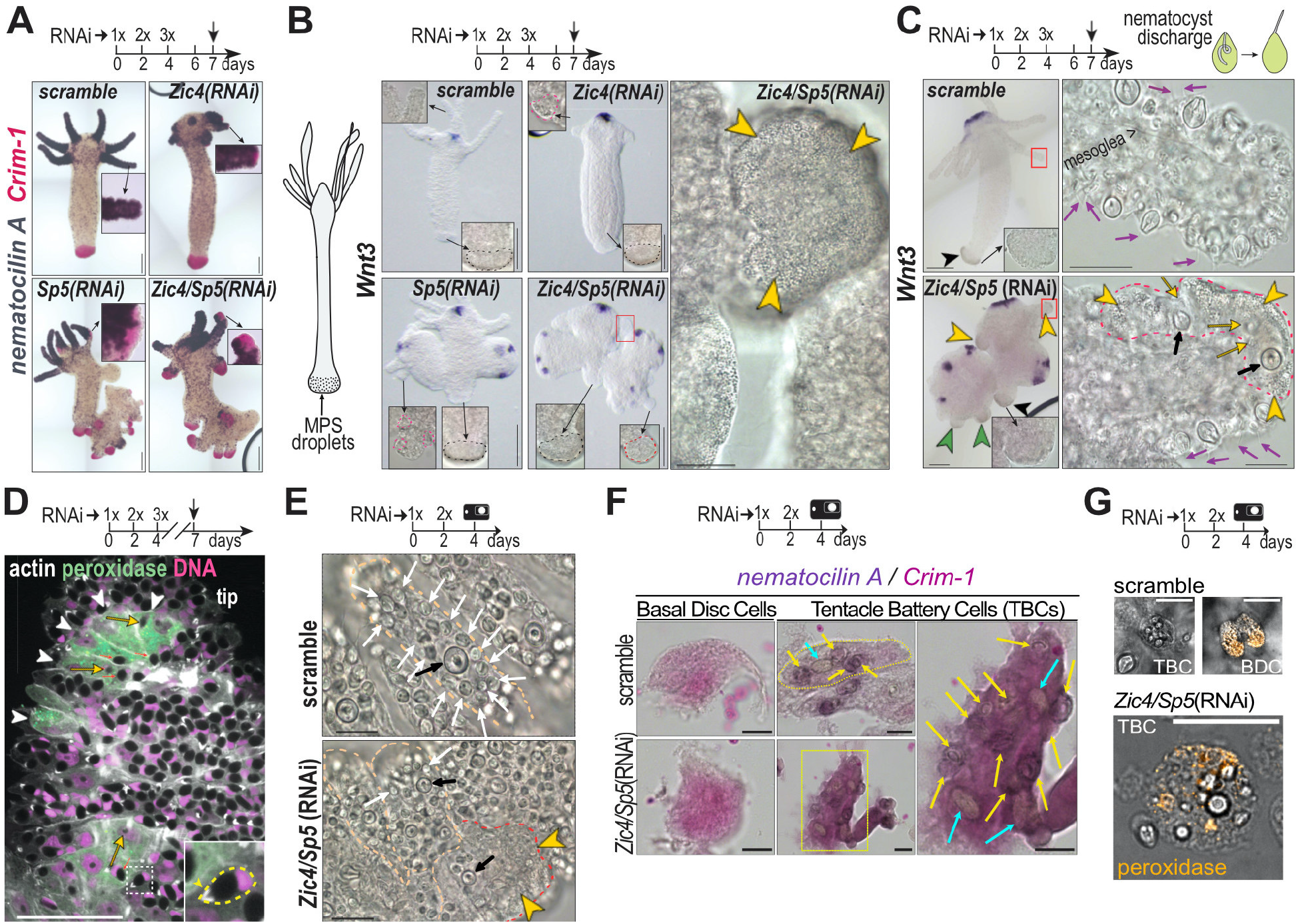
Dedifferentiation and basal differentiation of TBCs in *Hydra* knocked-down for *Zic4* and/or *Sp5*. **(A)** *Nematocilin-A* (purple) and *Crim-1* (pink) expression in *Hv_AEP2* knocked-down as indicated. **(B, C)** *Wnt3* expression and ectopic differentiation of basal tissue in *Hv_Basel* knocked-down as indicated. Black outline: original basal discs; red outline: ectopic basal tissue; red squares: enlarged transforming regions; yellow arrowheads: basal-like cells in tentacles characterized by abundant mucopolysaccharide (MPS) droplets; green arrowheads: basal-like cells in ectopic axial structures; purple arrows: discharged nematocysts; yellow arrows: degenerating SNs; black arrows: stenoteles. **(D)** *Zic4/Sp5*(RNAi) tentacle containing TBCs with peroxidase activity (white arrowheads). Inset: nematocyte with moon-shape nucleus and an actin-dense V-shape structure (white) at the nematocyst apex (arrowhead). Yellow arrows point to fuzzy nematocysts. **(E)** Disorganized *Zic4/Sp5*(RNAi) tentacle, containing both typical TBCs (yellow outline) with SN (white arrows) and stenoteles (black arrows) and transforming TBCs (red dotted line) with few SNs and a granular cytoplasm (yellow arrowheads). **(F)** Co-detection of *nematocilin-A* (purple) and *Crim-1* (pink) in macerates as indicated. Yellow square: enlarged cell; yellow arrows: SNs; blue arrows: LNs. **(G)** Peroxidase activity in BDCs and TBCs from basal and apical regions from *Hv_AEP2* animals treated as indicated, dissected and trypsin macerated.

We hypothesized that the formation of ectopic basal discs relies on the transdifferentiation of TBCs. As supporting evidence, we found TBCs, which exhibit a mixed identity, half tentacle, half basal disc three days post-EP3 with the typical basal mucus aspect in the close vicinity of degenerating nematocytes (**Figure 3C**, **Supplement Movie 1-3**). These TBCs actually contain lipid droplets with peroxidase activity, never observed in apical regions of scramble(RNAi) animals (**Figure 3D**). Four days later, *Zic4/Sp5*(RNAi) animals harbor tentacles fully transformed into basal discs characterized by an intense peroxidase activity (**Figure Supplement 24**). This transformation occurs already after two exposures to *Zic4/Sp5* siRNAs, as evidenced by tentacles containing two days post-EP2 TBCs with degenerated nematocytes, a typical granular cytoplasm (**Figure 3E, Figure Supplement 25**, **Supplement Movie 4-5**) and peroxidase+ droplets (**Figure Supplement 26**). We confirmed the early start of this transdifferentiation event by identifying in apical tissues macerated two days post-EP2 isolated TBCs that contain nematocytes expressing *nematocilin-A* together with the BDC marker *Crim-1* (**Figure 3F, Figure Supplement 27**), or isolated TBCs that contain droplets with peroxidase activity (**Figure 3G, Figure Supplement 28**). These results confirm the transdifferentiation process, i.e. TBCs that transiently co-express tentacle and basal disc markers.

To further investigate gene-expression changes that occur in tentacles of animals knocked-down for *Zic4* and/or *Sp5*, we used proximal and basal tentacle tissue for RNA-seq analysis. When projected into a PCA space, generated from the positional sequencing of different body parts, the overall identity of the RNAi tentacles samples seems to shift only marginally from the location of the intact tentacles (**Figure 4A, Figure Supplement 29, Dataset Supplement 3-4)**. However, when the identity map is constructed with epithelia-specific genes, we noted a clear shift from tentacle towards basal identity in all RNAi samples. All investigated conditions show an increase in the expression of basal markers, combined with a decrease of tentacle marker genes (**Figure 4B, Dataset Supplement 4**), with the strongest modulations in the tentacles of *Zic4/Sp5*(RNAi) animals but also in ectopic structures of *Sp5*(RNAi) animals. These results, which confirm the basal transformation of TBCs, suggest a key role for Zic4 as we recorded the highest level of *Zic4* silencing after *Zic4/Sp5*(RNAi) when compared to *Sp5*(RNAi) or *Zic4*(RNAi) (**Figure Supplement 30**).

**Figure 4.**
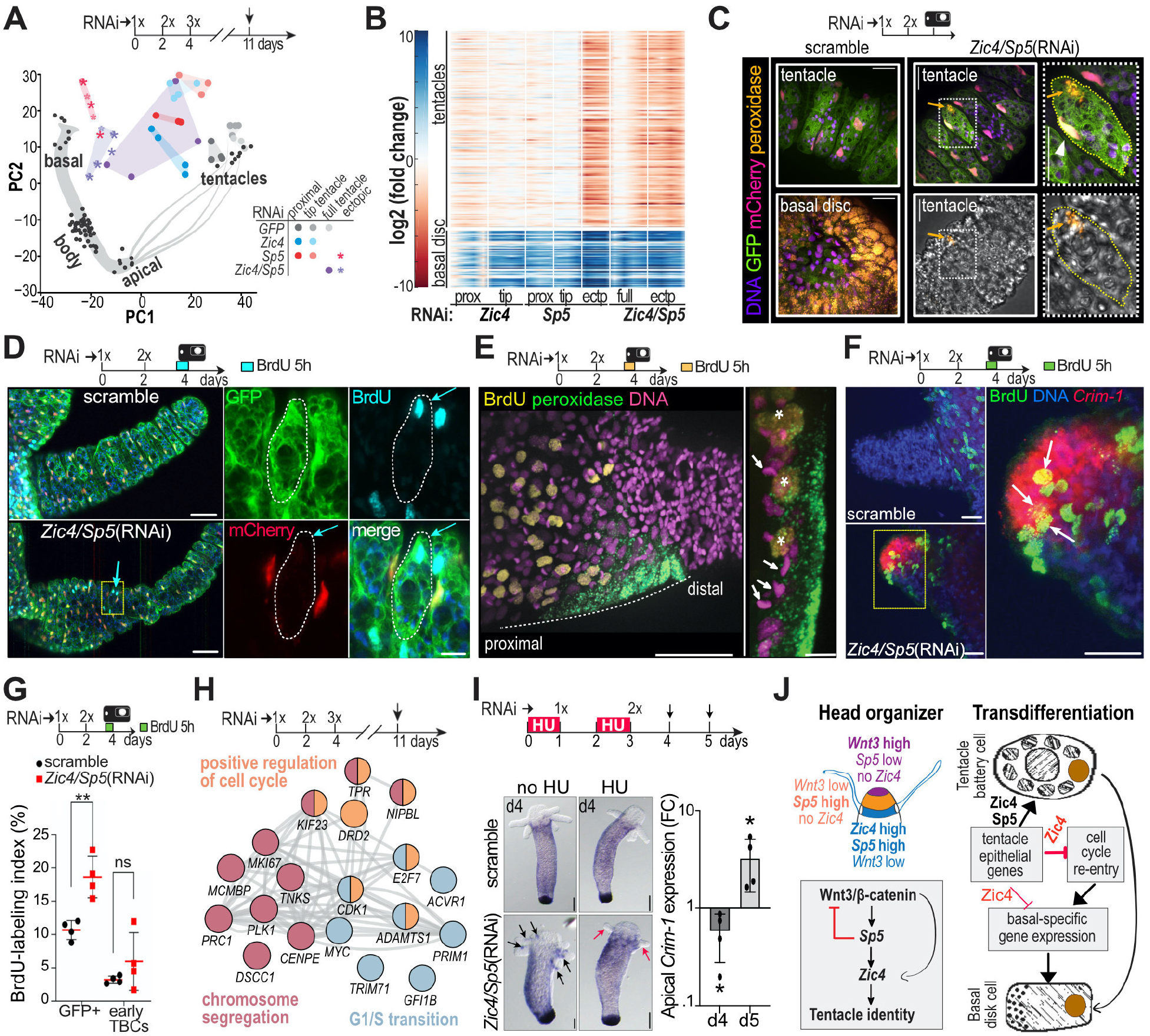
Transdifferentiating tentacle battery cells re-enter the cell cycle. **(A)** RNA-seq performed on dissected tentacles as indicated and projection into a PCA-space, generated by positional RNA-seq. **(B)** Heatmap showing differential expression of tentacle/basal disc markers. Full: whole tentacles, Prox, tip: proximal/distal tentacle regions, ectp: ectopic structures. **(C)** Detection of peroxidase (arrows) in FUCCI-eGFP animals. **(D)** BrdU+ TBC (blue arrows) enlarged on the right (white outline). **(E)** Peroxidase+ droplets (green) in BrdU+ TBC; right: enlarged section along the dotted line, asterisks: BrdU+ peroxidase+ cells (yellow), white arrows: nematocyte nuclei. **(F)** *Crim-1+* BrdU+ tentacle cells (white arrows). **(G)** BrdU-labeling index in apical epithelial cells (GFP+) and early TBCs. **(H)** Subnetwork of cell cycle genes upregulated in tentacles across all RNAi conditions (depicted in panel B); color-coding indicates GO terms, edges genetic/physical interactions, co-localization and/or co-expression. **(I)** Left: Ectopic *Crim-1* (black arrows) no longer detected after HU-treatment (red arrows). Right: *Crim-1* qPCR. **(J)** Left: Head organizer defined by *Wnt3/ Sp5/Zic4* domains and signaling cascade for tentacle identity. Right: Transdifferentiation model. P-values: **≤ 0.01, *< 0.05. Error bars indicate SD. Scale bars: 25 μm (C, F), 50 μm (D) 10 μm for enlargement, 50 μm (E) 20 μm for enlargement, 200 μm (I).

Next, we investigated whether cells that show signs of transdifferentiation also re-enter the cell cycle. First, we analyzed tentacles of FUCCI-eGFP *Hydra* knocked-down twice for *Zic4/Sp5* and we noted in cells that contain peroxidase+ droplets, a change from red/pink to yellow nuclear fluorescence, suggesting a progression through S/G2 phase (**Figure 4C, Figure Supplement 31**). To confirm cell cycle re-entry, we performed a 5-hour BrdU labeling of FUCCI-eGFP *Hydra* knocked-down for *Zic4/Sp5.* Two days post-EP2, we identified in tentacles BrdU+ nuclei among TBCs (**Figure 4D, Figure Supplement 32**), peroxidase+ droplets in BrdU+ TBCs (**Figure 4E, Figure Supplement 33B**), *Crim-1* expression in BrdU+ TBCs (**Figure 4F, Figure Supplement 33A**) and finally an increase in the number of BrdU+ apical epithelial cells quantified on macerated tissues or on whole-mounts (**Figure 4G, Figure Supplement 34**) but no increase in the number of dividing cells **(Figure Supplement 35**). We found that after four days of *Zic4/Sp5* silencing, the typical organization of TBCs becomes disrupted in the proximal part of tentacles of several animals, with a sharp boundary with the distal part that remain well organized (**Figure 4E, Figure Supplement 34**).

To collect evidence on the molecular mechanism that possibly drives cell-cycle re-entry, we analyzed the GO term of the genes consistently upregulated when *Zic4* and/or *Sp5* are knocked-down and we identified biological functions linked to cell cycle, i.e. G1/S transition, positive regulation of the cell cycle, chromosome segregation (**Figure 4H, Dataset Supplement 5**), confirming that epithelial cell cycling is enhanced upon *Zic4* and/or *Sp5* silencing. To test whether cell cycling is necessary for tentacle transdifferentiation, we inhibited DNA synthesis with two pulses of Hydroxyurea (HU). We observed that ectopic *Crim-1* spots no longer form in tentacles of *Zic4/*Sp5(RNAi) animals, while apical *Crim-1* expression is transiently abrogated, in agreement with the transient HU-induced blockade of cell cycle progression (Buzgariu et al. 2018) (**Figure 4I, Figure Supplement 36**). Taken together, these data indicate that upon *Zic4/Sp5* silencing, TBCs do not maintain their typical tentacle character, re-enter the cell cycle but do not divide, and fully transform within less than seven days into basal disc cells.

The results obtained in intact and regenerating animals indicate that distinct levels of Wnt/β-catenin signaling, Sp5 and Zic4 activity define three regions in the apical region and trigger the formation of two distinct structures, the hypostome and the tentacle zone. In the hypostome Wnt/β-catenin signaling is high, while Zic4 and Sp5 are low. The tentacle zone is characterized by low Wnt/β-catenin signaling and high *Sp5*, inducing *Zic4* expression and thus TBC differentiation and tentacle formation. The enhanced transdifferentiation phenotype obtained in animals submitted to the double *Zic4*/*Sp5* knockdown is coherent with an epistatic relationship between *Sp5* and *Zic4*. We conclude that *Zic4* acts as a master regulator to control the choice between two epidermal cell fates, maintenance or differentiation of battery cells when *Zic4* is high, differentiation of basal mucous cells when *Zic4* is low. Therefore, we suspect Zic4 to repress the basal disc cell status, which appears as a default state of epithelial epidermal differentiation, possibly reflecting an ancestral differentiation fate of pre-bilaterian multifunctional epithelial stem cells. In the sea anemone *Nematostella*, several Zic genes are expressed during tentacle formation in distinct tentacle cell types (Layden et al. 2010), suggesting a function in tentacle formation possibly shared among cnidarians.

We also present evidence for epithelial transdifferentiation in *Hydra*, with a model where low levels of expression of the Zic4 and Sp5 transcription factors lead to cell fate change. In such context, clustered tentacle battery cells first dedifferentiate, namely lose *ks1* expression, no longer maintain embedded nematocytes as evidenced by the loss of *NemA* expression, while the typical tentacle architecture gets disorganized as the large extra-cellular space between TBCs disappears. In parallel, these TBCs re-enter the cell cycle without dividing and re-differentiate into basal disc cells that express *Crim-1*, differentiate droplets with peroxidase activity at the apical pole first and rapidly form a compact mass (**Figure 4J, Figure Supplement 37**). As TBCs that express both tentacle and basal markers could be identified, we concluded that this process fulfills the criteria of transdifferentiation (Lambert et al. 2021). In addition, cell division was not observed among tentacle BrdU-positive cells, and mitotic activity not detected in tentacles, therefore, we conclude that cells that re-enter S-phase undergo endoreplication, a process observed in developmental as well as injury and stress contexts, associated with regeneration (Lee et al. 2009)(Nandakumar et al. 2021).

Zinc finger transcription factors (ZF-TF) play a crucial role in cell fate stability, the ectopic expression of a single ZF-TF (Zfp521) being able to promote the conversion of human fibroblasts to neural stem cells *ex vivo* (Shahbazi et al. 2016). Among them, the Zic family members act as versatile multifunctional proteins, classical DNA-binding proteins that regulate the expression of the ascidian *Brachyury* or the mammalian *Oct4* and *Nanog* via their promoter sequences. *Zic* genes also act as transcriptional co-activators via their protein-protein interaction with a variety of transcription factors such as Gli, TCF, Smad, Pax, Cdx, SRF as well as chromatin remodeling factors contributing to enhancer functions (Hatayama and Aruga 2018). Of interest, in vertebrates, Zic proteins inhibit Wnt/β-catenin signaling (Fujimi et al. 2012), compete with Sp transcription factors on promoter sequences (Yang et al. 2000), or associate with Geminin in cell cycle regulation (Sankar et al. 2016). The current identification of a Zic4/Wnt-dependent GRN that acts as a switch regulating the fate of epithelial cells, opens the way for deeper investigation of this GRN in evolutionary-distant eumetazoan phyla.

## Supporting information

Supplement VOGG

## Acknowledgments

We thank all members of the Tsiairis and Galliot lab for discussions, Denis Duboule for comments on the manuscript. The authors thank the iGE3 Genomic Platform for RNA-seq library preparation and sequencing, the FMI Genomics facility and Sebastien Smallwood for RNA sequencing experiments, the FMI Imaging facility for support with microscopy, and Iskra Katic for *Hydra* electroporations.

## Author contributions

Conceptualization: MCV, JF, SVi, BG, CDT; Methodology: MCV, JF, WB, BG, CDT; Investigation: MCV, JF, WB, CP, PP, PGLS, CN, CD, CR, SVr; Visualization: MCV, JF, WB, PGLS, LB, BG, CDT; Funding acquisition: BG, CDT; Supervision: BG, CDT; Writing – original draft: MCV; Writing – review & editing: BG, MCV, JF, WB, PGLS, SVi, CDT

## Data availability

The RNA-seq data have been deposited in the GEO database under the accession codes GSE190110 and GSE191177. All other raw data can be found in Dataset Supplement 6.

