## Supplement VOGG for "The transcription factor Zic4 acts as a transdifferentiation switch"

### Supplementary Figures

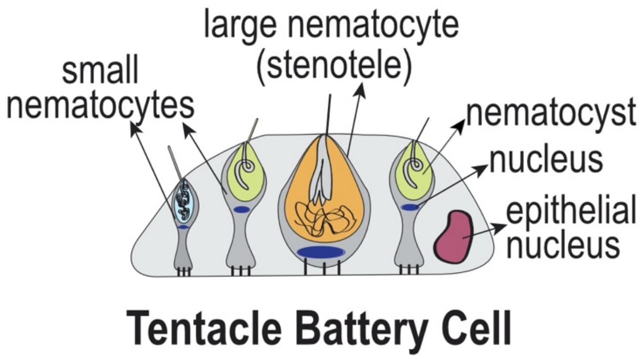


**Figure Supplement 1. Schematic view of a Tentacle Battery Cell (TBC)**
Tentacle Battery Cell (TBC) are epidermal epithelial cells located exclusively in the tentacles, which embed nematocytes. Early TBCs contain up to 9 nematocytes whereas mature TBCs contain up to 20 small nematocytes centered around one or two large nematocytes. Nematocytes or stinging cells are mechano-sensory cells equipped with a cnidocil, acting as a sensory organ, and a venom capsule named nematocyst that upon stimulation of the cnidocil discharges its venom through the discharge of the tubule (Hufnagel et al. 1985; Holstein 2012). Several types of nematocytes can be identified, large ones that contain a large nematocyst named *stenotele* with a prominent stylet apparatus used upon discharge to pierce the epidermis/cuticle of the preys; small nematocytes that are characterized by the type of nematocyst they contain: either small *desmonemes* with a tightly coiled tubule, or *holotrichous and spineless atrichous isorhizas*. Note the compressed nucleus (blue), basal to the nematocyst.

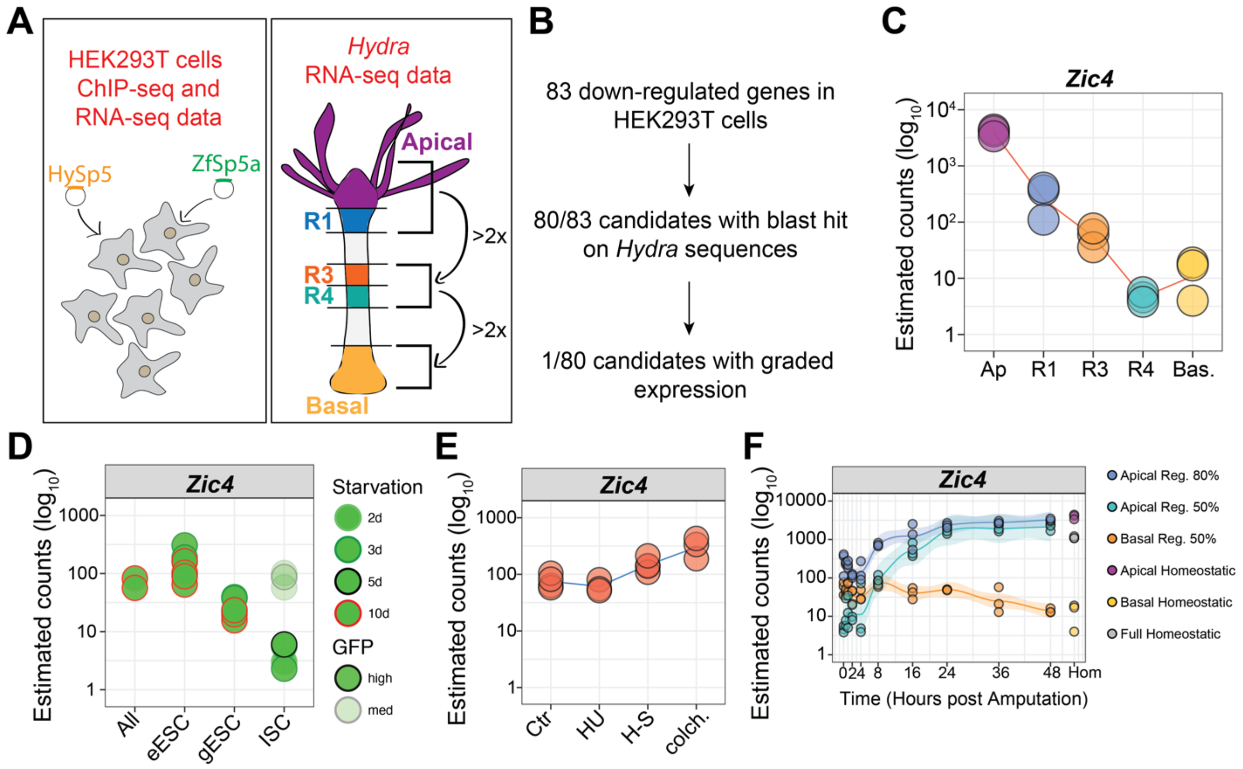


**Figure Supplement 2. Characterization of Zic4 as a putative Sp5 target gene in *Hydra* vulgaris (Hv)**

(**A-B**) Procedure to identify *Sp5* target genes. Previous work had found 83 human genes down-regulated in HEK293T cells overexpressing *HySp5* and *ZfSp5a*, identified as putative direct *Sp5* target genes (Vogg et al. 2019). 80 *Hydra* orthologs (Dataset S1) were retrieved from the *Hv*_Jussy transcriptome with an E-value < $10^{-3}$ (HydrAtlas.unige.ch) (Wenger et al. 2019). To identify genes showing an apical-to-basal graded expression, we compared for each gene the RNA-seq expression values obtained in tissues taken at five positions along the body axis (apical -Ap-, regions R1, R3, R4, basal -Bas-) (HydrAtlas.unige.ch) and selected genes with a minimum 2-fold decrease between the apical region+R1 vs R3+R4 and a 2-fold decrease between the R3+R4 vs the basal region. The transcription factor *Zic4* was the unique gene to fulfill this criterion, see Dataset S1. (**C-F**) RNA-seq expression profiles of *Zic4* (*seq19466_loc08275*) in intact and regenerating *Hv*_Jussy animals (C, F), in *Hv_AEP2* stem cell populations (D, *c11132_g1_i01*), after the elimination of interstitial cells (E).

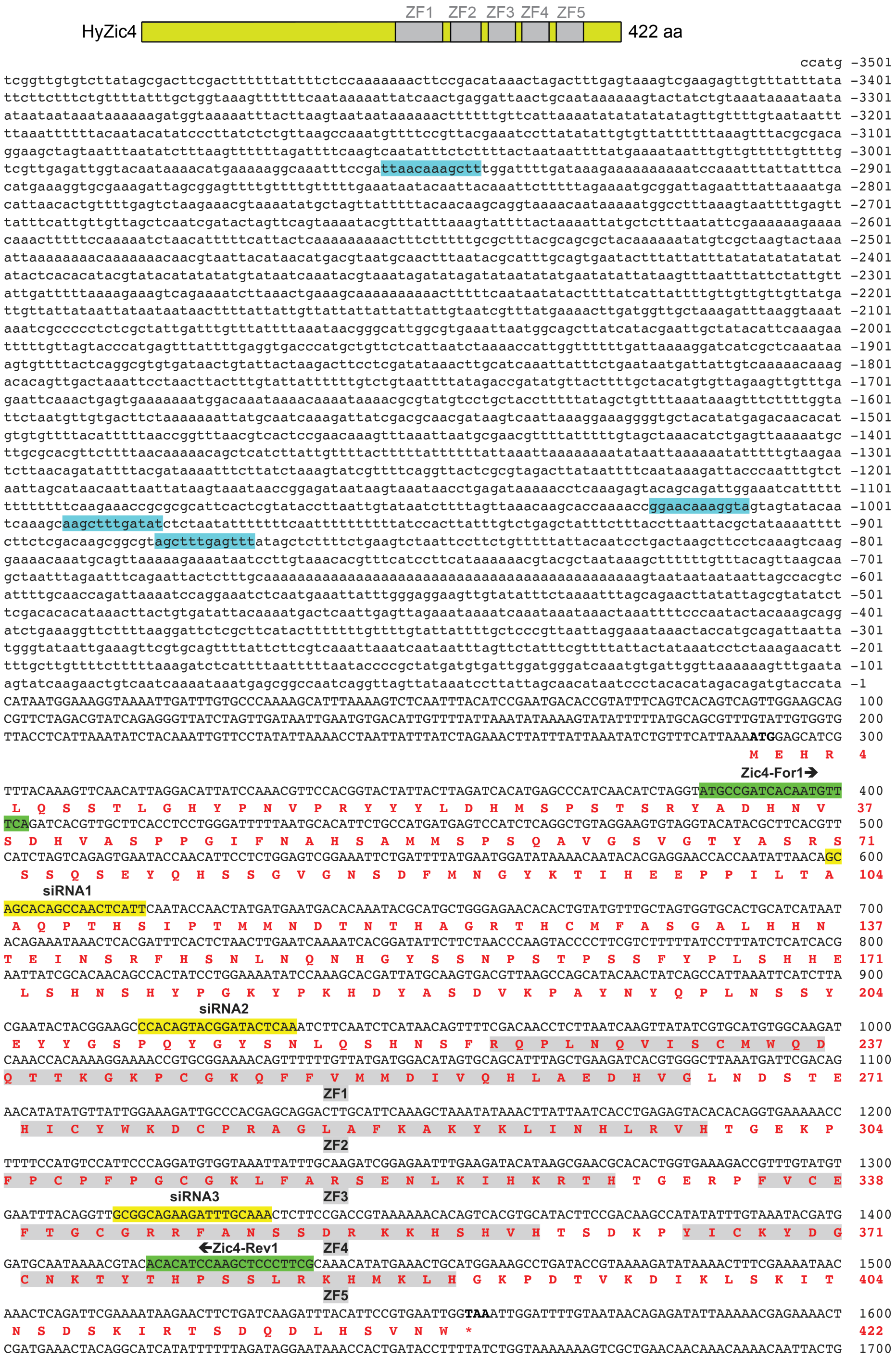


**Figure Supplement 3.** ***Hydra Zic4*** **sequence and structure of the** ***Zic4*** **transcription factor**

The *Zic4* genomic sequence is written lowercase and the coding sequence uppercase. The five zinc finger domains (ZF1-5) are highlighted in grey, TCF binding sites in blue, siRNA sequences in yellow and the primers used for cloning into pGEM-T Easy and subsequent riboprobe generation in green.

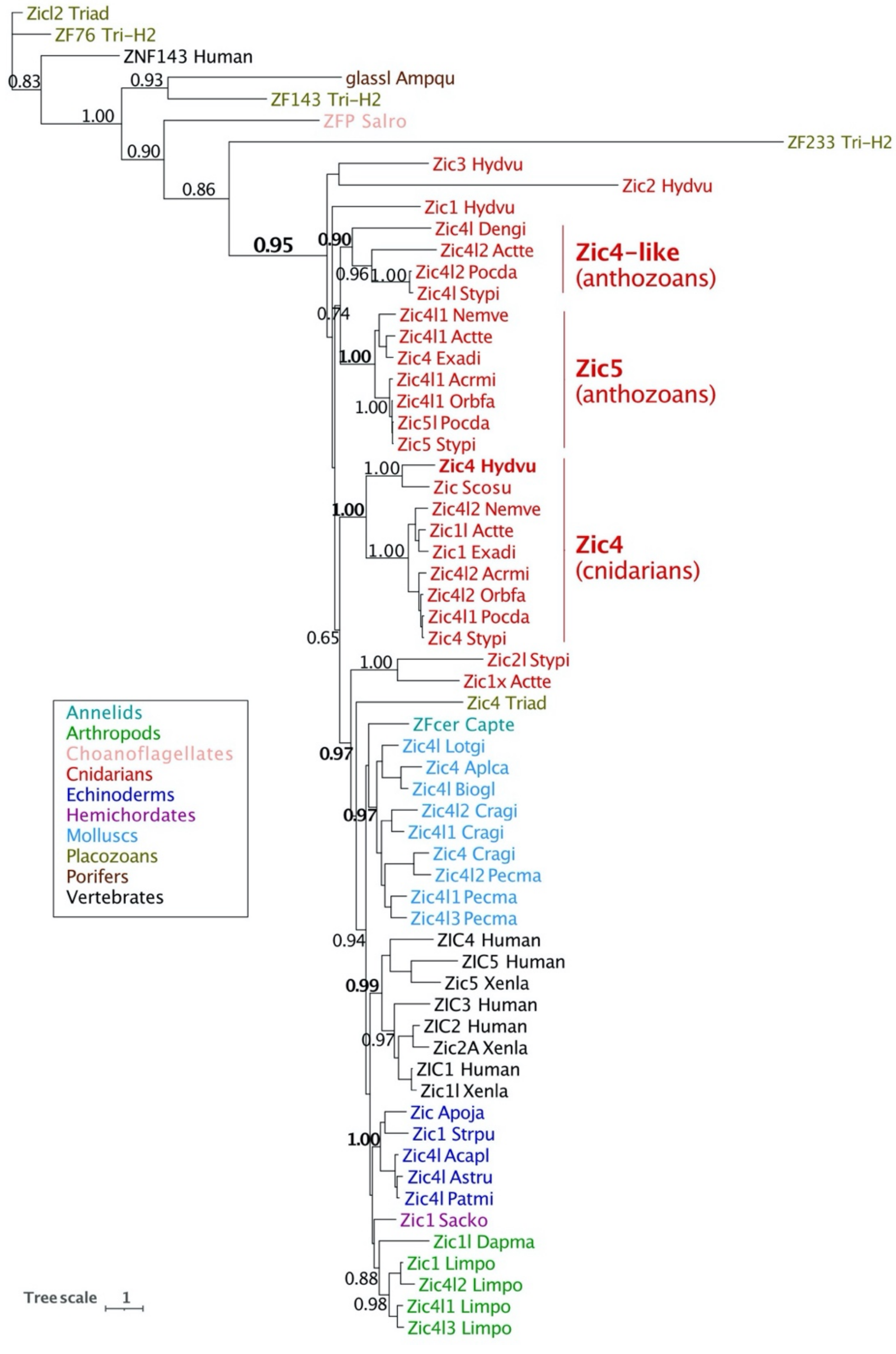


**Figure Supplement 4. Phylogenetic analysis of the Zinc-Finger Zic gene family**

The color code indicates the different phyla as written in the legend. *Hydra* expresses four distinct Zic-related genes (*Zic1, Zic2, Zic3, Zic4*) and only the Zic4 family contains sequences from both hydrozoan and anthozoan species. In contrast, the Zic4l and Zic5 families appear anthozoan-specific. In bilaterians, the diversification of the Zic gene families has arisen several times independently. Note the absence of Zic-related sequences in pre-eumetazoan phyla. For the species code and the accession numbers of each gene, see Table S1.

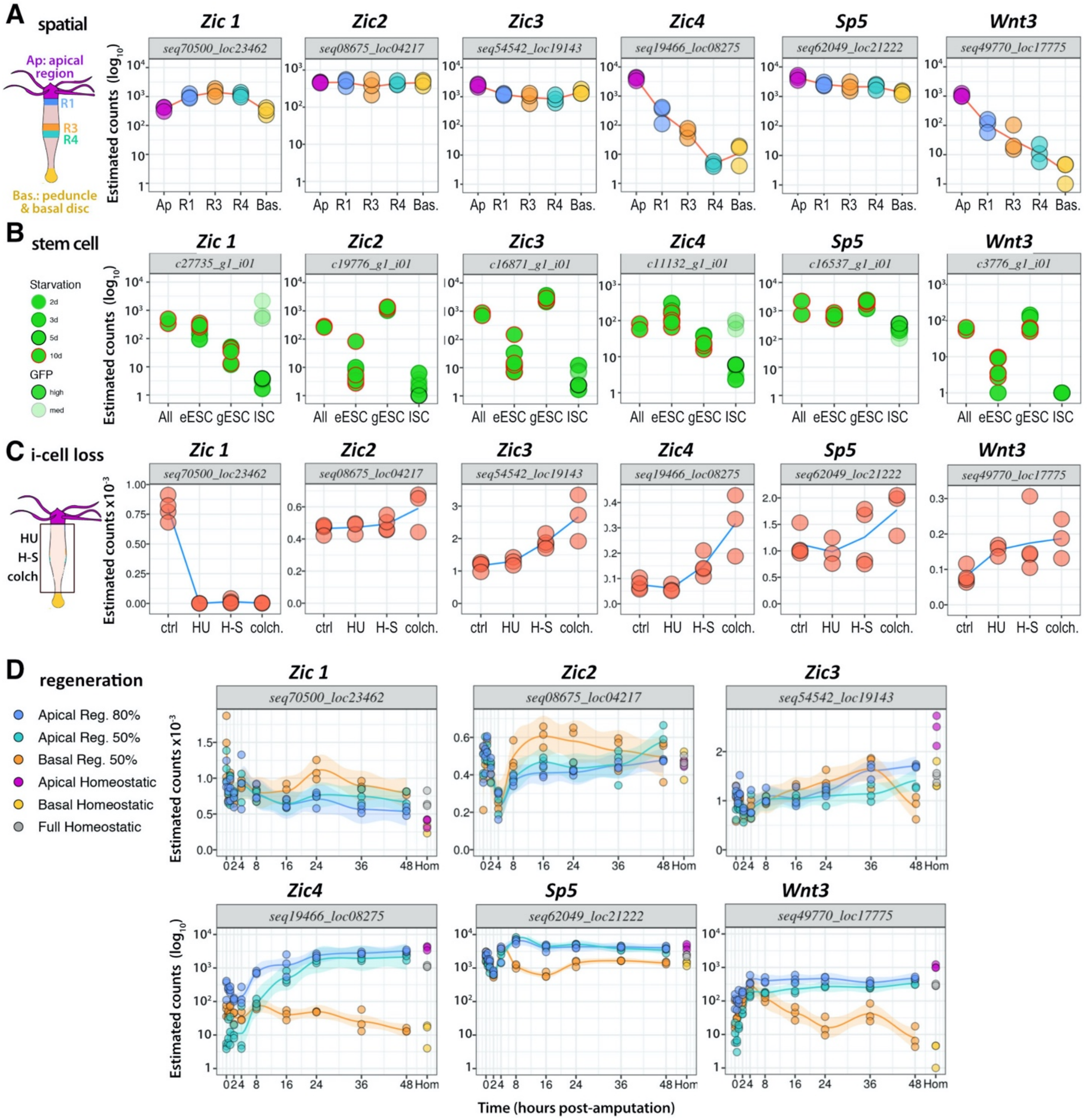


**Figure Supplement 5. Comparative analysis of the expression profiles of the *Zic*, *Sp5* and *Wnt3* genes in *Hydra***

(**A**) Expression profiles were measured by RNA-seq at five distinct locations along the body axis as indicated on the left. (**B**) Expression profiles were measured by RNA-seq in each of the three stem cell populations present in transgenic animals from the *H. vulgaris AEP2* strain that constitutively expresses GFP either in the epidermal Epithelial Stem Cells (eESC), gastrodermal ESC (gESC), or in the Interstitial Stem Cells (ISC). Animals were starved for variable periods of time, the central gastric regions dissected, and the GFP-expressing cells sorted by flow cytometry and subsequently used for RNA-seq analysis. All: RNA-seq analysis performed on unsorted cells from the gastric region. (**C**) RNA-seq analyses performed on *H. vulgaris* animals from the thermosensitive strain *sf-1* taken seven days after the exposure either to hydroxyurea (HU), heat-shock (H-S) or to colchicine (colch.) as described in (Wenger et al. 2016). Note that except *Zic1*, all *Zic, Sp5* and *Wnt3* genes tend to get up-regulated after the elimination of the interstitial cells (i-cell loss). (**D**) RNA-seq analyses performed on apical or basal-regenerating tips dissected at nine distinct time-points of regeneration, either after decapitation (Apical Regeneration 80% - blue) or after mid-gastric bisection (Apical Regeneration 50% - green- Basal Regeneration 50% - orange-). Hom: homeostatic values indicated on the right of the graph. In (A) and (D), RNA-seq analyses were performed on *H. vulgaris* animals from the Jussy strain. All experiments were repeated at three different periods of the year. For technical details, see in (Wenger et al. 2016; Wenger et al. 2019). In (C) and (D) upper row, the y scale is linear as the modulations remain moderate, whereas in all other panels the y scale is logarithmic (log10). Note the apical-to-basal graded distribution of the *Zic4, Sp5* and *Wnt3* transcripts; *Zic4* displays the strongest quantitative modulations along the axis as well as during regeneration.

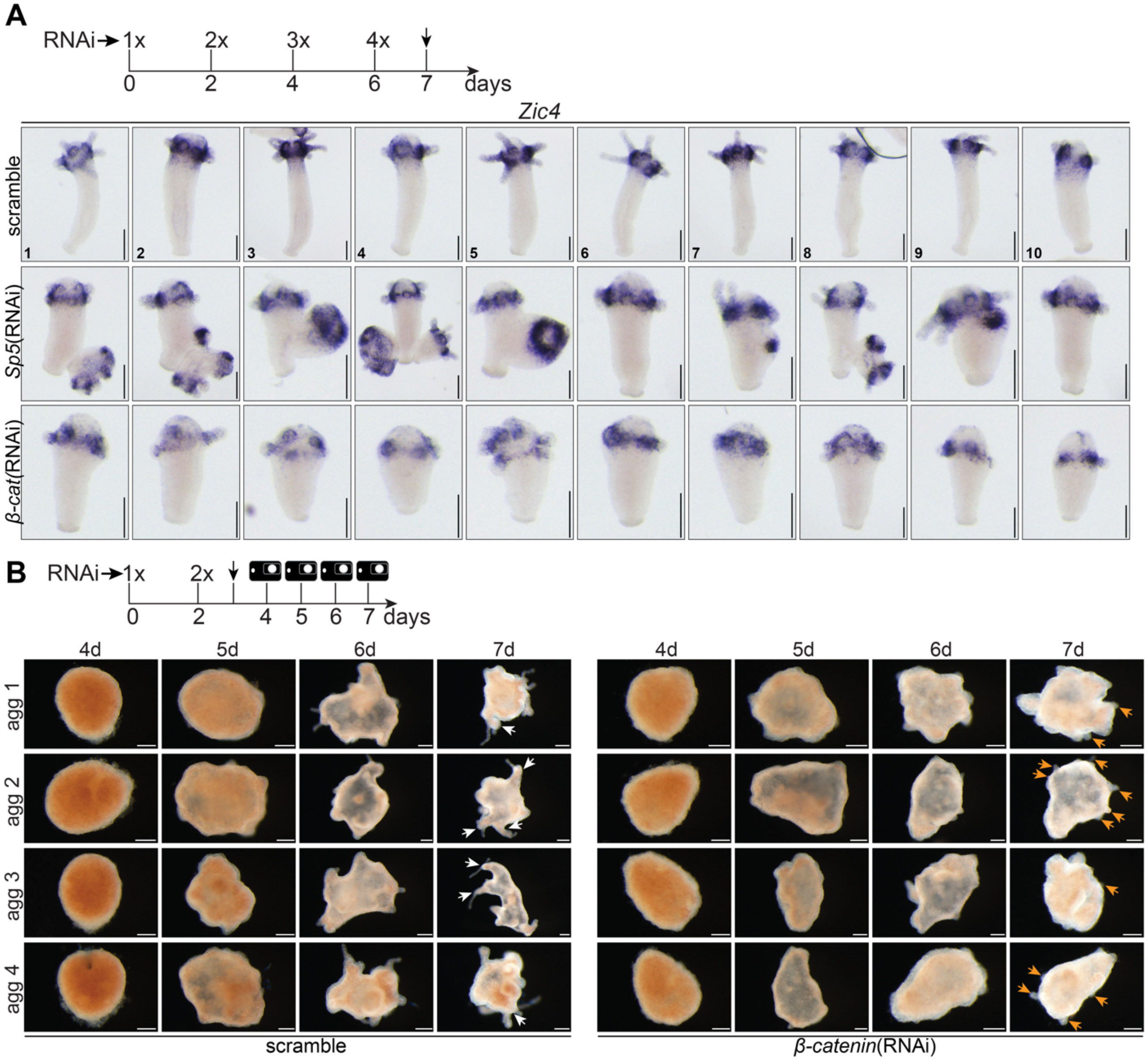

**Figure Supplement 6. Impact of *Sp5*(RNAi) and *β-catenin*(RNAi) on *Zic4* expression and apical differentiation**

**(A)** Intact *Hv_Basel* were electroporated four times every other day (RNAi1-4) and fixed one day after the last electroporation for *in situ* hybridization. Shown are ten representative animals of an experiment performed in triplicate. **(B)** Knockdown of *β-catenin* in reaggregation studies. Intact *Hm-105* were electroporated twice (RNAi1, RNAi2) with a scramble siRNA or a mix of *β-catenin* siRNAs, dissociated one day after RNAi2 to obtain a cell suspension and immediately reaggregated. Reaggregates (agg1-4) were imaged live on day 4, 5, 6 and 7 after reaggregation. Note that *β-catenin*(RNAi) reaggregates form only a few tentacles (orange arrows) while axis are clearly visible in scramble aggregates (white arrows). Scale bars: 200 µm.

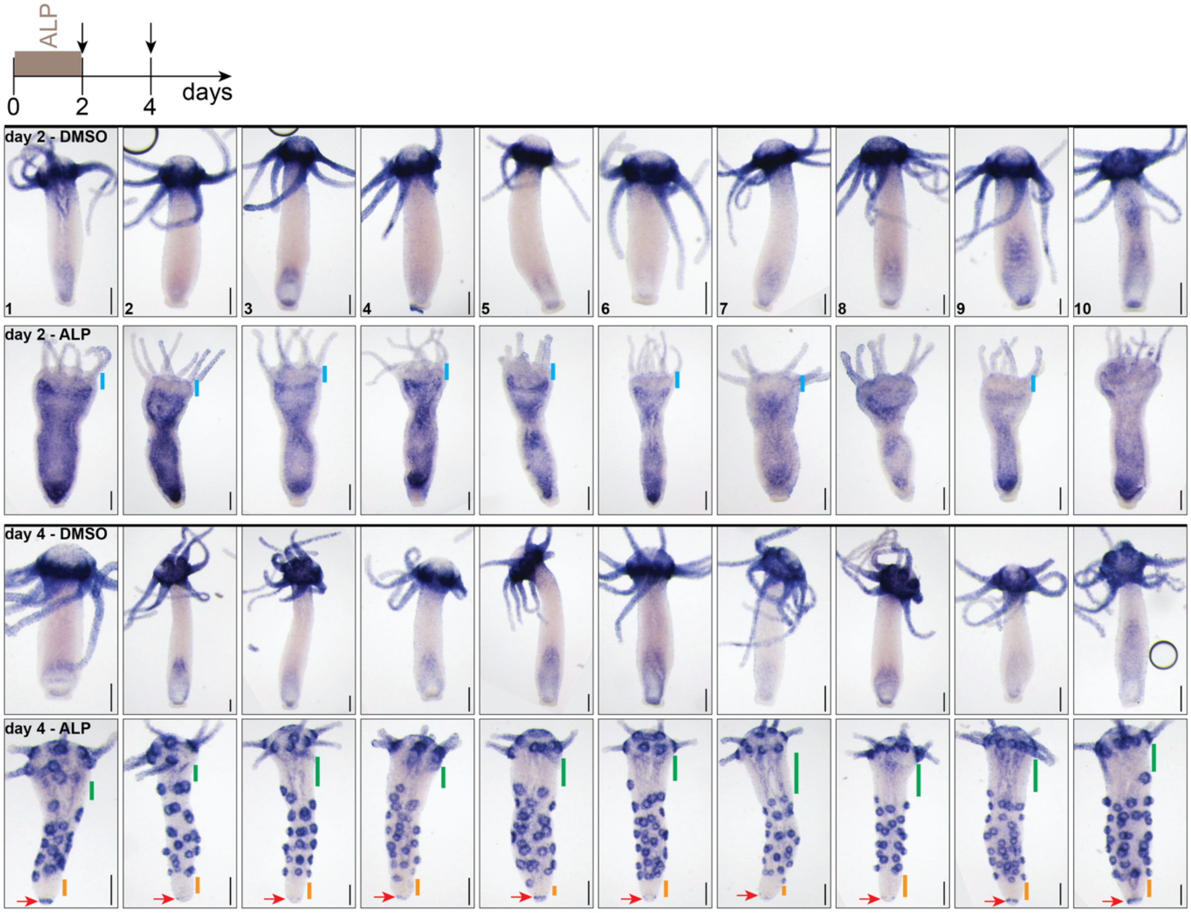


**Figure Supplement 7. *Zic4* expression in Alsterpaullone (ALP) treated animals**

Intact *Hv_Basel* exposed for 2 days to Alsterpaullone (ALP) were fixed on day 2 and day 4 for in *situ* hybridization. Immediately after ALP treatment (day 2), *Zic4* expression is detected in the body column but decreased in the head (blue line). Two days later (day 4), *Zic4* is expressed in multiple rings along the central body column but absent from the most upper and lower regions of the body column (green and orange lines). Note the expression of *Zic4* in the basal disc (red arrows) in ALP-treated animals. Ten representative animals of an experiment performed in duplicate are shown. Scale bars: 200 µm.

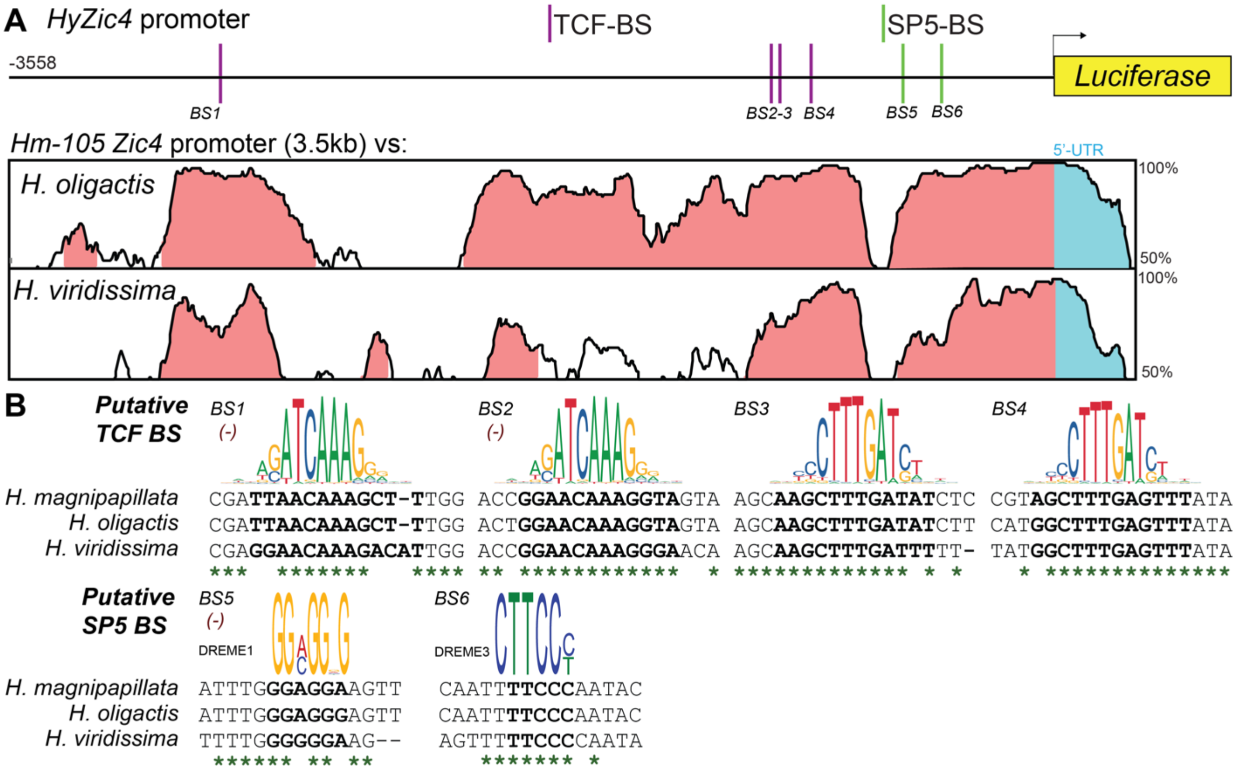


**Figure Supplement 8. Mapping of putative TCF and Sp5 binding sites in the *Zic4* promoter**

(**A**) Map of the 3'558 bp genomic region encompassing the *Zic4* promoter of *H. magnipapillata* (*Hm-105*) and phylogenetic footprinting plot comparing this region to the corresponding region in the *H. oligactis* and *H. viridissima* genomes. Evolutionarily conserved modules (at least 70% base-pair identity over a 100 bp sliding window) are shown in pink in the Vista alignment plot. Putative TCF-BS were numbered according to their 5'-3' location relative to the *Zic4* TSS. (**B)** Sequence alignment displaying evolutionary conservation across the three *Hydra* species analyzed for putative TCF and Sp5-BS highlighted in (A). For Sp5-BS prediction, the DREME matrix corresponding to the identified motif is indicated (see Material and Methods). The sequence corresponding to the predicted TCF-BS is highlighted in bold. The PMW logo for the TCF and Sp5 binding sites are displayed above each alignment. In those cases in which the putative BS mapped in the reverse DNA strad (-), the reverse-complement version of the matrix was used for graphical visualisation. Nucleotide positions conserved across the three species are marked by asterisks.

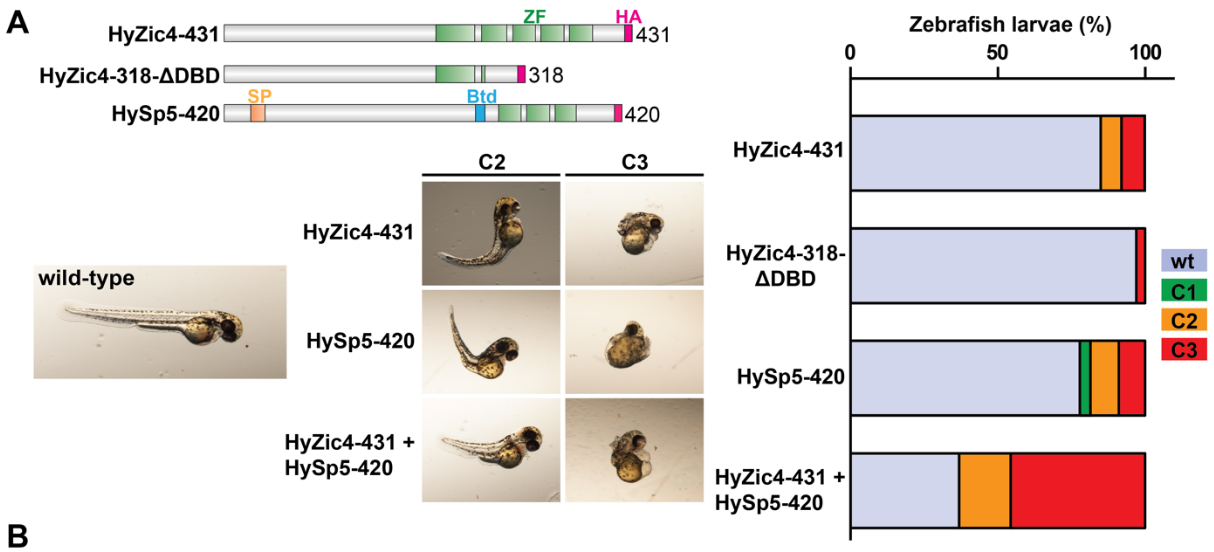


**B**

| mRNA | [conc] pg | N | †% (n) | Sn | wt % (n) | class1 % (n) | class2 % (n) | class3 % (n) | Total % of abnormal embryos |
|---|---|---|---|---|---|---|---|---|---|
| **HyZic4-431** | 10 | 132 | 4.6% (6) | 126 | **84.9%** (107) | **0.0%** | **7.1%** (9) | **7.9%** (10) | **15.0%** |
| **HyZic4-318-ΔDBD** | 10 | 109 | 7.3% (8) | 101 | **97.0%** (98) | **0.0%** | **0.0%** | **3.0%** (3) | **3.0%** |
| **HySp5-420** | 400 | 115 | 1.7% (2) | 113 | **77.9%** (88) | **3.5%** (4) | **9.7%** (11) | **8.9%** (10) | **22.1%** |
| **HyZic4-431 + HySp5-420** | 10+400 | 132 | 22.0% (29) | 103 | **36.9%** (38) | **0.0%** | **17.5%** (18) | **45.6%** (47) | **63.1%** |

**Figure Supplement 9. Overexpression of *Hydra Zic4* in zebrafish embryos**

(**A**) Full length *HyZic4* (HyZic4-431), *HyZic4* lacking part of the DNA binding domain (HyZic4-318-ΔDBD) and full length *Sp5* (HySp5-420) mRNAs were injected into zebrafish embryos and screened for morphological defects on day 2 post-fertilization. The phenotypes were scored into three classes: straight axis with eye phenotype (C1); curly axis with eye phenotype (C2); underdeveloped axis, curly tail, and eye phenotype (C3). The bar graph represents the percentage of zebrafish larvae with morphological defects. Note that the co-injection of *HyZic4* and *HySp5* mRNAs increases the phenotypic penetrance. (**B**) Table summarizing the phenotypic analysis. N = number of injected embryos; †%(n) = percentage (number) of dead embryos; Sn = number of survived embryos. These data correspond to an experiment performed in duplicate.

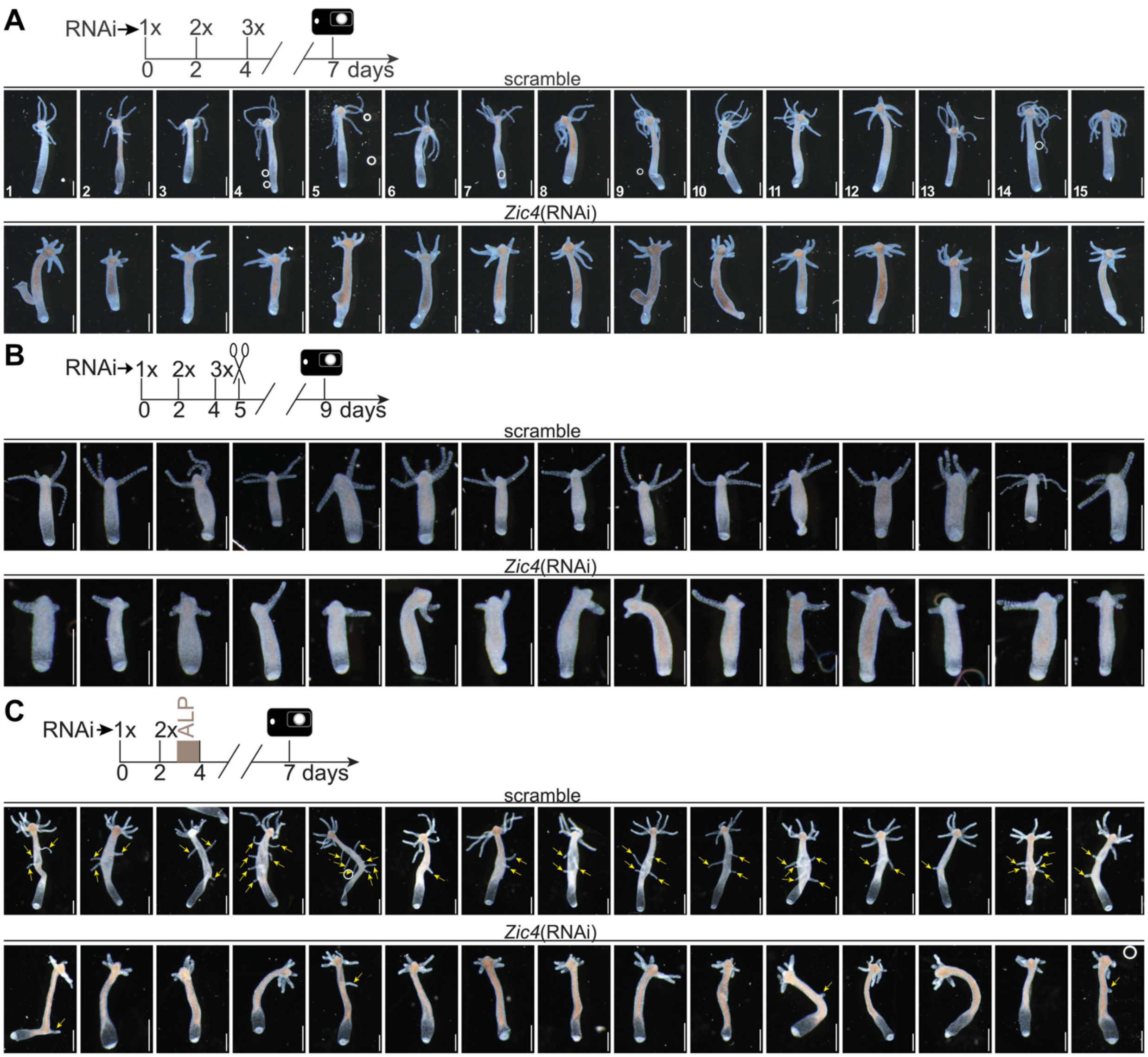


**Figure Supplement 10. *Zic4* is required for tentacle formation**

Tentacle defect observed after silencing *Zic4* in intact (**A**), head regenerating (**B**) and Alsterpaullone (ALP) treated (**C**) *Hv_Basel*. Apical regeneration was induced by amputating the animals 50% body length (mid-gastric bisection). Electroporations were performed every other day (RNAi1, RNAi2, RNAi3 in A-B; RNAi1, RNAi2 in C) and animals fixed and imaged on the indicated days. Yellow arrows point towards ectopic tentacles. Note the reduction in ectopic tentacle formation after *Zic4*(RNAi). Shown are fifteen representative animals of an experiment performed in triplicate (n = 15 each). Scale bars: 500 µm.

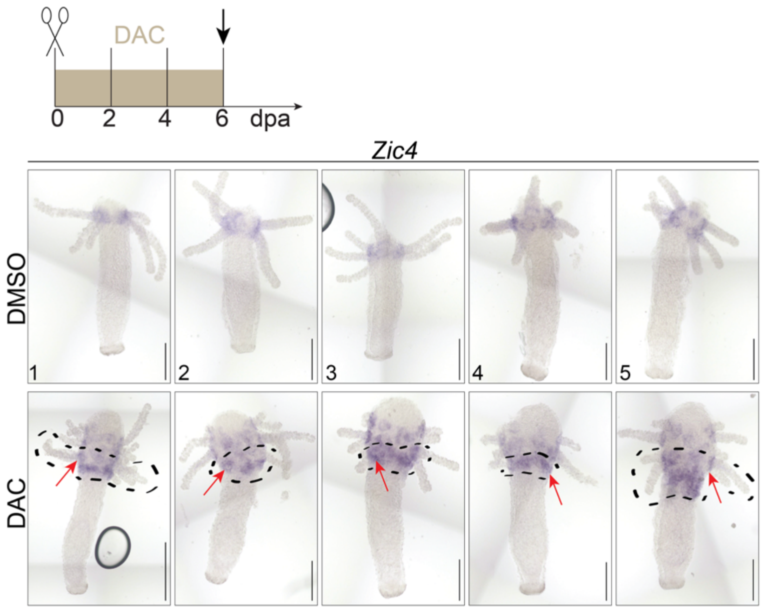


**Figure Supplement 11.** ***Zic4*** **expression in DAC-treated animals**

*Zic4* expression in *Hv_Basel* animals having regenerated their head while treated or not with DAC-2-25 for 6 days after mid-gastric bisection. *Zic4* is expressed at the tentacle base (red arrows). Note that DAC-2-25 causes the formation of a second tentacle ring (encircled in black) (Glauber et al. 2013). Shown are five representative animals of an experiment performed in duplicate. Scale bars: 250 µm.

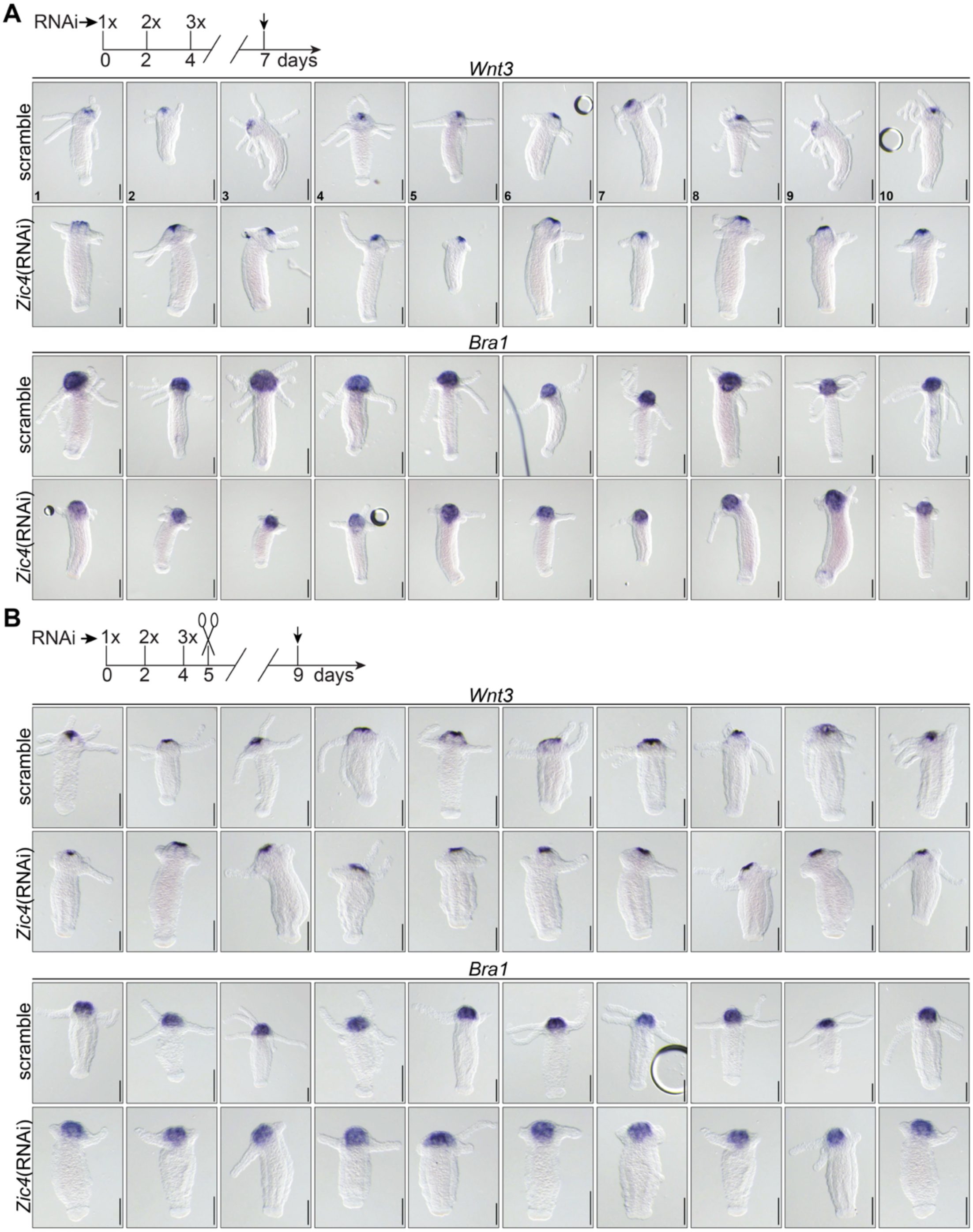


**Figure Supplement 12. *Wnt3* and *Bra1* expression in intact and apical-regenerating *Zic4*(RNAi) *Hydra***

Intact (**A**) and 4-day apical-regenerating (**B**) *Hv_Basel* animals were electroporated three times with a scramble siRNA or *Zic4* siRNAs and processed for *in situ* hybridization on day 7 and day 9, respectively to detect *Wnt3* and *Bra1* expression. All experiments were performed in triplicates (n = 10 each). Note that *Zic4*(RNAi) does not affect the expression of *Wnt3* and *Bra1*. Scale bars: 200 µm.

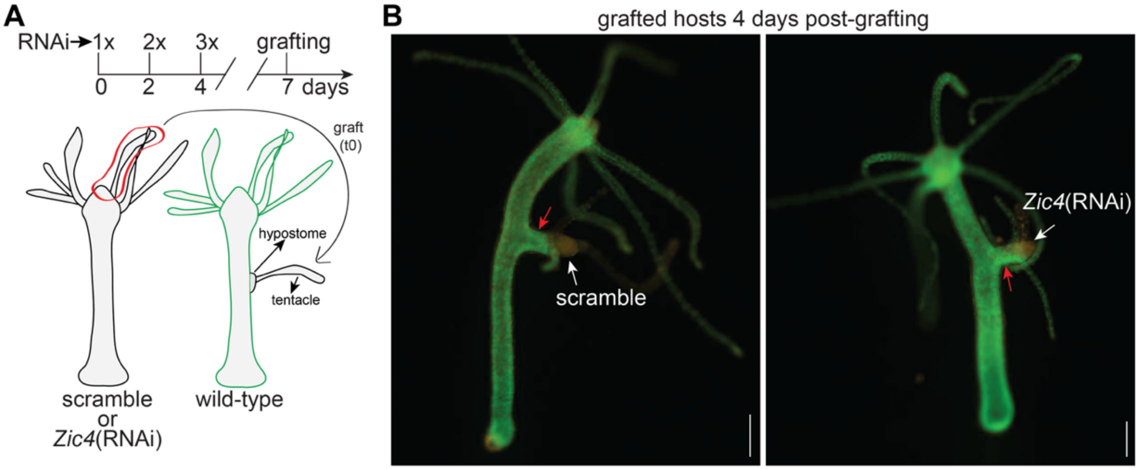


**Figure Supplement 13. Knocking-down *Zic4* expression does not affect head organizer activity** (**A**) Experimental overview. Intact non-transgenic *Hv_AEP2* animals were electroporated every other day with a scramble siRNA or a mix of *Zic4* siRNAs. On day 7, a piece of hypostomal tissue was grafted onto the body column of a GFP+ host. The hypostomal tissue was grafted together with a tentacle that does not possess organizer activity and that served as a marker of the graft. (**B**) All hosts grew ectopic axes (red arrows) following the transplantation of grafts (white arrows) derived either from scramble or from *Zic4*(RNAi) animals. Shown are representative animals of an experiment performed in duplicate (n1 = 3 animals per condition; n2 = 4 animals per condition). Scale bars: 250 µm.

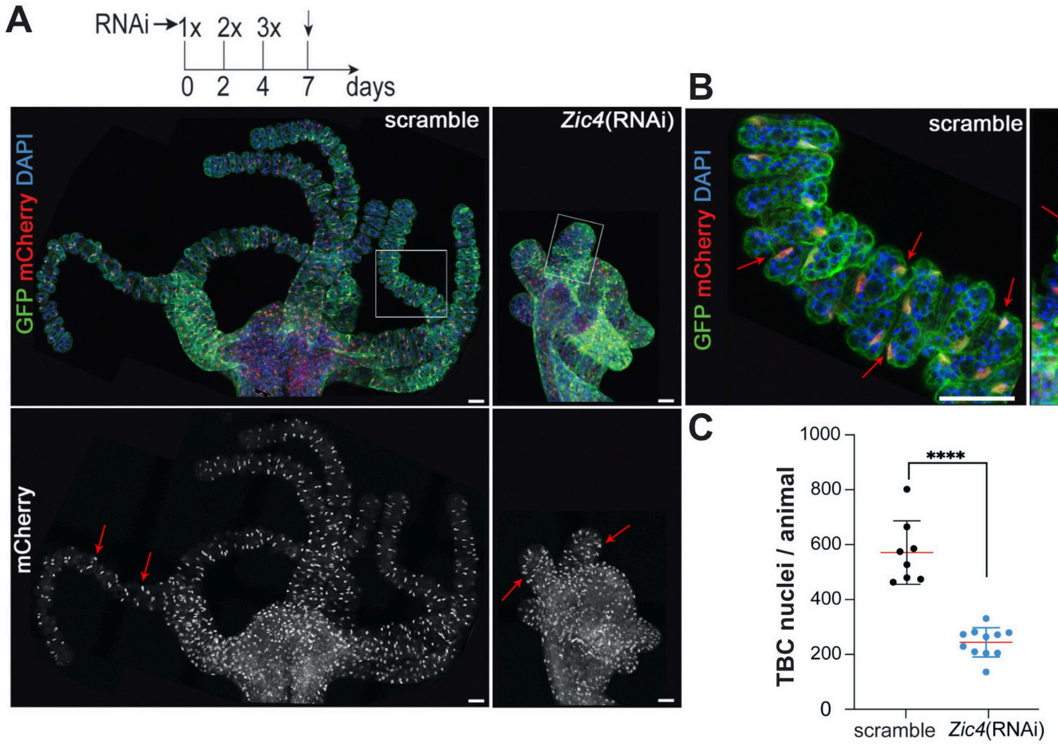


**Figure Supplement 14. Reduction in the TBC number in tentacles of *Zic4*(RNAi) FUCCI animals**

Transgenic FUCCI-eGFP *Hydra* that constitutively express Geminin-GFP (green, cytoplasmic) and CTD-mCherry (red, nuclear) in the epidermal epithelial cells shown here three days after the third electroporation (EP3) with scramble or *Zic4* siRNAs, immunostained and DAPI-stained nuclei. **(A)** Images of scramble and *Zic4*(RNAi) *Hydra* that represent the maximal projection of a Z-stack covering 30 µm. The red arrows point to epidermal nuclei mCherry+ in the tentacles. The white rectangles indicate the enlarged regions presented in **(B)** for both scramble and *Zic4*(RNAi) animals. Scale bars: 50 µm. **(C)** To quantify the number of TBCs, the number of mCherry+ nuclei was counted in each tentacle and the total number of epidermal nuclei/animal was represented. Each data point represents one animal. Error bars indicate SD. Statistical p-value: ****≤ 0.0001.

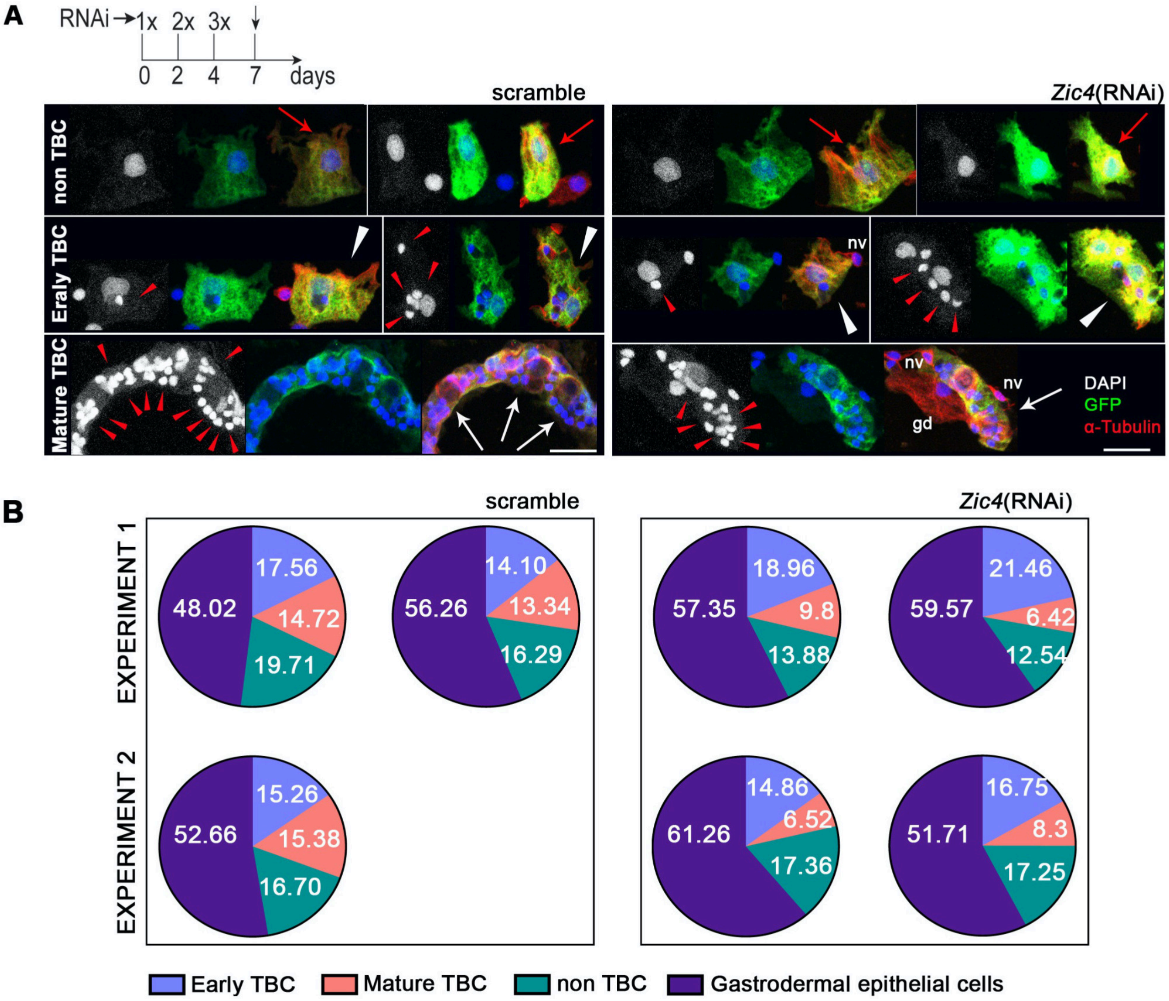


**Figure Supplement 15. Proportion of apical epithelial cell types in *Zic4*(RNAi) FUCCI animals**
FUCCI-eGFP *Hydra* that constitutively express Geminin-GFP and CTD-mCherry in the epidermal epithelial cells were electroporated three times with a scramble siRNA or a mix of *Zic4* siRNAs. Three days post-EP3, the apical region was dissected, macerated and cells were immunostained for GFP (green) and alpha-tubulin (red). The GFP+ epidermal epithelial cells were classified in three categories as described in (Dübel et al. 1987): non-TBC epidermal epithelial cells, early TBCs that contain up to 9 nematocytes, mature TBCs that contain 10-20 nematocytes. Cells from each category from scramble and *Zic4*(RNAi) animals **(A)** are shown with red arrows pointing to non-TBC epidermal epithelial cells, white arrowheads to early TBCs, white arrows to mature TBCs and red arrowheads to nematocytes. Abbreviations: nv: nerve cells; ic: interstitial cells; gd: gastrodermal epithelial cell. Scale bars: 25 µm. **(B)** Charts representing the relative proportion of the different apical epithelial cell types in scramble and *Zic4*(RNAi) animals. Two experiments with one or two replicates per condition were performed and GFP+ cells from the entire slide were manually characterized and quantified. Few GFP+ nuclei coming from broken epidermal cells were included in the non-TBC epidermal epithelial cell category, and few GFP-negative TBCs with a typical morphology were also included in the early or mature TBC categories.

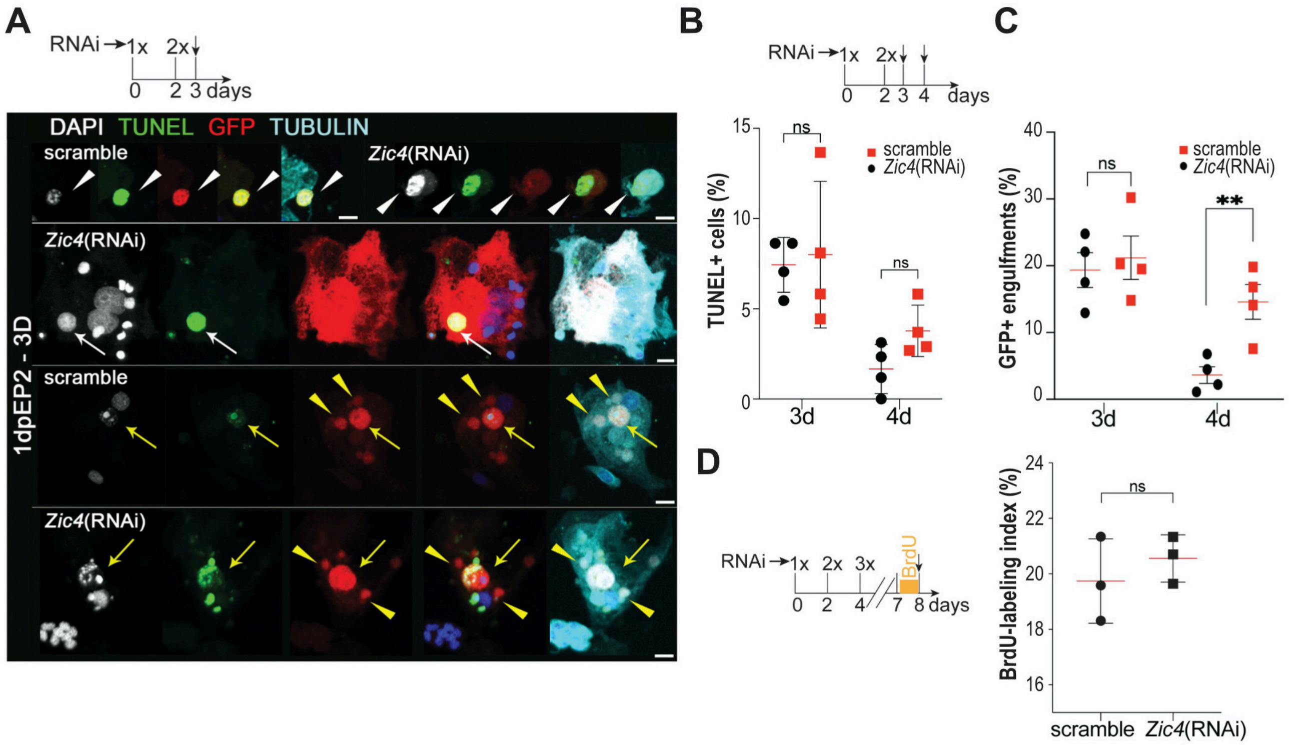


**Figure Supplement 16. Epithelial cell death and epithelial cell cycling in *Zic4*(RNAi) animals**
(**A-C**) The apical region of transgenic FUCCI-eGFP *Hydra* electroporated twice with scramble or *Zic4* siRNAs were dissected on day 1 and day 2 after EP2, macerated to detect the apoptotic cells by TUNEL. (A) Representative apoptotic cells from scramble and *Zic4*(RNAi) animals detected by TUNEL (green), immunostained for GFP (red) and tubulin (cyan) and nuclei DAPI-labeled (white). White arrowheads point to late apoptotic cells that have lost their cytoplasm; white arrows point to an early apoptotic GFP+ cell in a cluster of three epidermal GFP+ cells from a *Zic4*(RNAi) animal. Gastrodermal epithelial cells have engulfed GFP+ apoptotic bodies (yellow arrows) and/or GFP+ vacuoles (yellow arrowheads). Scale bars: 10 µm. (B-C) The graphs show the percentage of TUNEL+ cells among the epidermal epithelial cells (B) and the percentage of epithelial cells containing GFP+/TUNEL+ and/or GFP+ debris (C). Data were obtained from two independent experiments, each with two replicates. At least, 900 cells were counted for each condition. Statistical p-value: **≤ 0.01. Errors bars indicate SD. (**D**) *Hv_Basel* animals electroporated three times with scramble or *Zic4* siRNAs were incubated in BrdU for 16 hours three days after EP3. On day 8, the body column was dissected, macerated and the BrdU-labeling index calculated as the percentage of BrdU+ epithelial cells among epithelial cells. Each data point represents one independent biological experiment where at least 900 cells were counted.

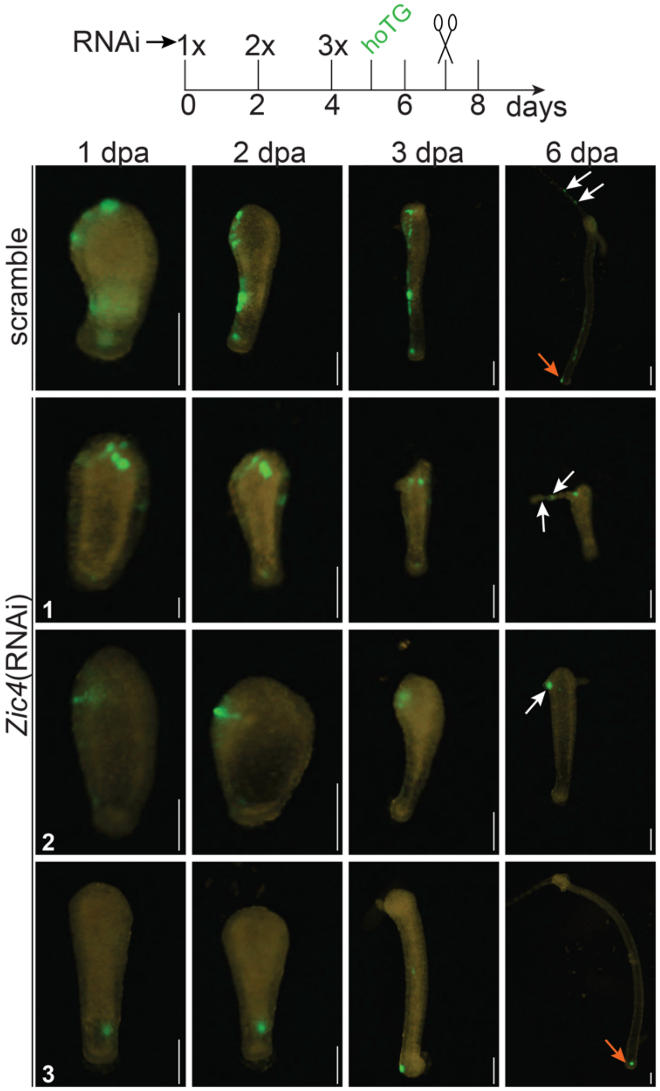


**Figure Supplement 17. Displacement of epithelial cells in *Zic4*(RNAi) animals**

Intact *Hv_Basel* were electroporated three times with a scramble siRNA or a mix of *Zic4* siRNAs and on day 5 electroporated with the hoTG plasmid (Wittlieb et al. 2006). The animals were cut 50% body length (mid-gastric bisection) on day 7 and imaged 1, 2, 3 and 6 days post amputation (dpa). Note that the displacement of epithelial cells is not deficient in *Zic4*(RNAi) animals. White arrows point towards GFP+ cells that were displaced towards the tentacles/head and red arrows point towards GFP+ cells that were displaced towards the foot. Scale bars: 200 µm.

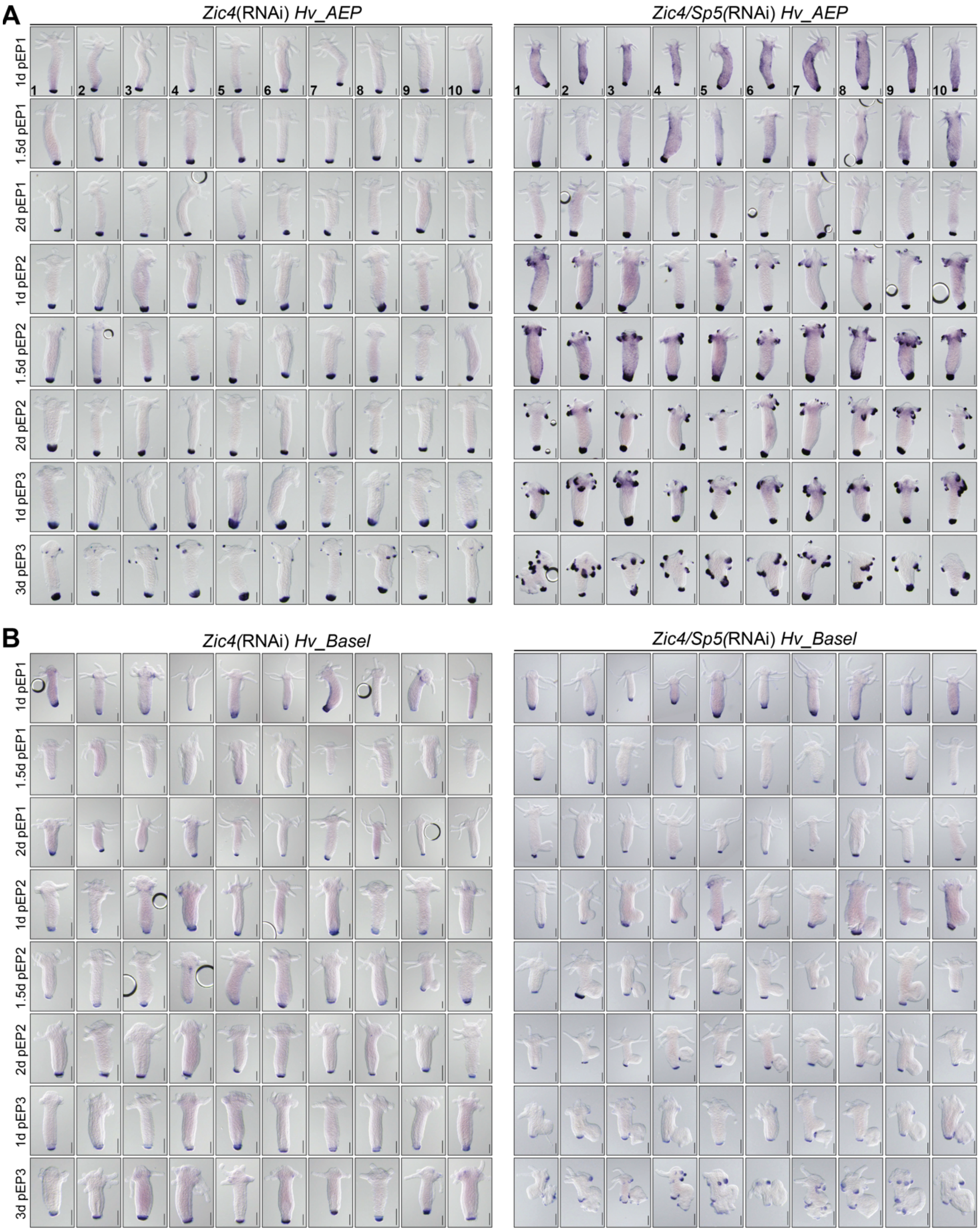


**Figure Supplement 18. Ectopic *Crim-1* expression in *Zic4*(RNAi) and *Zic4*/*Sp5*(RNAi) *Hv_AEP2* or *Hv_Basel* animals**

Intact *Hv_AEP2* (**A**) and *Hv_Basel* (**B**) animals electroporated (EP) three times with *Zic4* siRNAs or with a mixture of *Zic4* and *Sp5* siRNAs were fixed at indicated time points after EP1, EP2, EP3. 10 representative animals are shown for each condition. Scale bars: 200 µm.

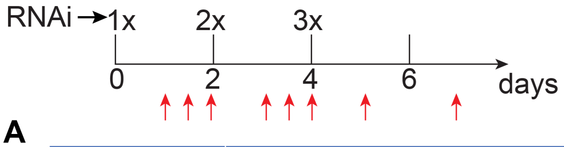


**A**

| | | Ectopic apical *Crim-1* in *Hv_AEP* | | | | | | Ectopic apical *Crim-1* in *Hv_Basel* | | | | | |
|---|---|---|---|---|---|---|---|---|---|---|---|---|---|
| conditions | | *Zic4*(RNAi) | | | *Zic4/Sp5*(RNAi) | | | *Zic4*(RNAi) | | | *Zic4/Sp5*(RNAi) | | |
| **RNAi1** | 1d | 6/20 | 30% | ± | 20/20 | 100% | + | 8/18 | 44.4% | + | 14/17 | 82.4% | + |
| | 1.5d | 1/18 | 5.6% | ± | 20/20 | 100% | + | 3/17 | 17.6% | + | 5/19 | 26.3% | ± |
| | 2d | 3/18 | 16.7% | ± | 15/19 | 79% | + | 4/18 | 22.2% | ± | 2/19 | 10.5% | ± |
| **RNAi2** | 1d | 7/20 | 35% | ± | 18/18 | 100% | ++ | 11/18 | 61.1% | + | 12/19 | 63.2% | + |
| | 1.5d | 6/17 | 35.3% | + | 15/15 | 100% | +++ | 5/16 | 31.3% | + | 3/14 | 21.4% | + |
| | 2d | 6/18 | 33.3% | + | 17/17 | 100% | +++ | 4/16 | 25% | + | 5/20 | 25% | + |
| **RNAi3** | 1d | 15/18 | 83.3% | + | 19/19 | 100% | +++ | 10/20 | 50% | + | 14/16 | 87.5% | + |
| | 3d | 17/19 | 89.5% | ++ | 20/20 | 100% | +++ | 24/28 | 85.7% | + | 20/20 | 100% | ++ |

**B**

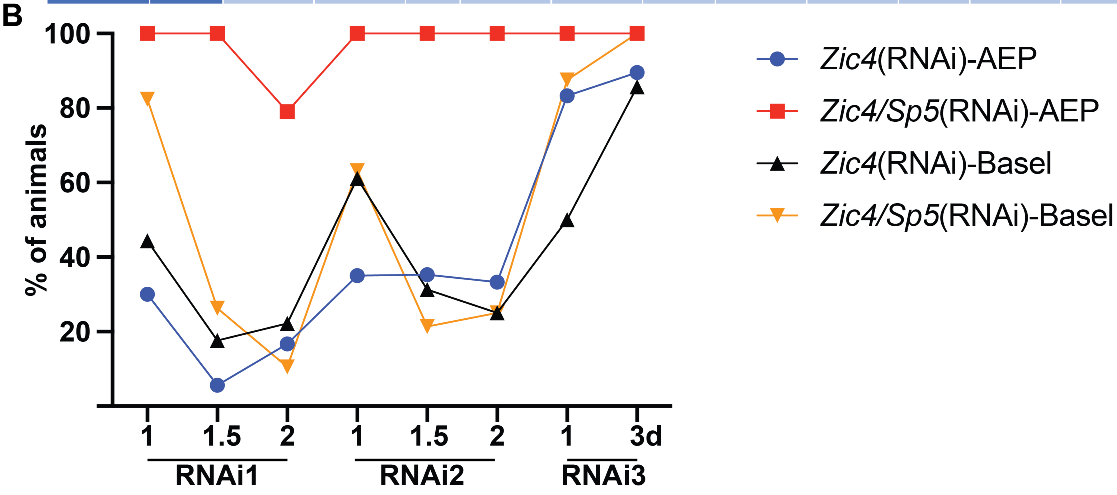


**Figure Supplement 19. Comparative quantitative analysis of apical *Crim-1* expression after *Zic4* or *Zic4/Sp5*(RNAi)**

(**A**) Number and percentage of *Hv_AEP2* or *Hv_Basel* animals showing ectopic *Crim-1* expression at 8 distinct time-points as depicted in Figure S18. The intensity of ectopic *Crim-1* expression is considered as homogenous for a given condition, corresponding to one of the five levels: absent (-); low (±); moderate (+); high (++); very high (+++). (**B**) Graphical representation of the data shown in panel (A). Note the transient ectopic *Crim-1* expression after RNAi1 and the higher number and intensity of ectopic *Crim-1* expression in *Hv_AEP2* after *Zic4/Sp5*(RNAi) when compared to *Hv_Basel*. The total number of animals was obtained from two distinct experiments with n= 7-10 for each condition in each experiment.

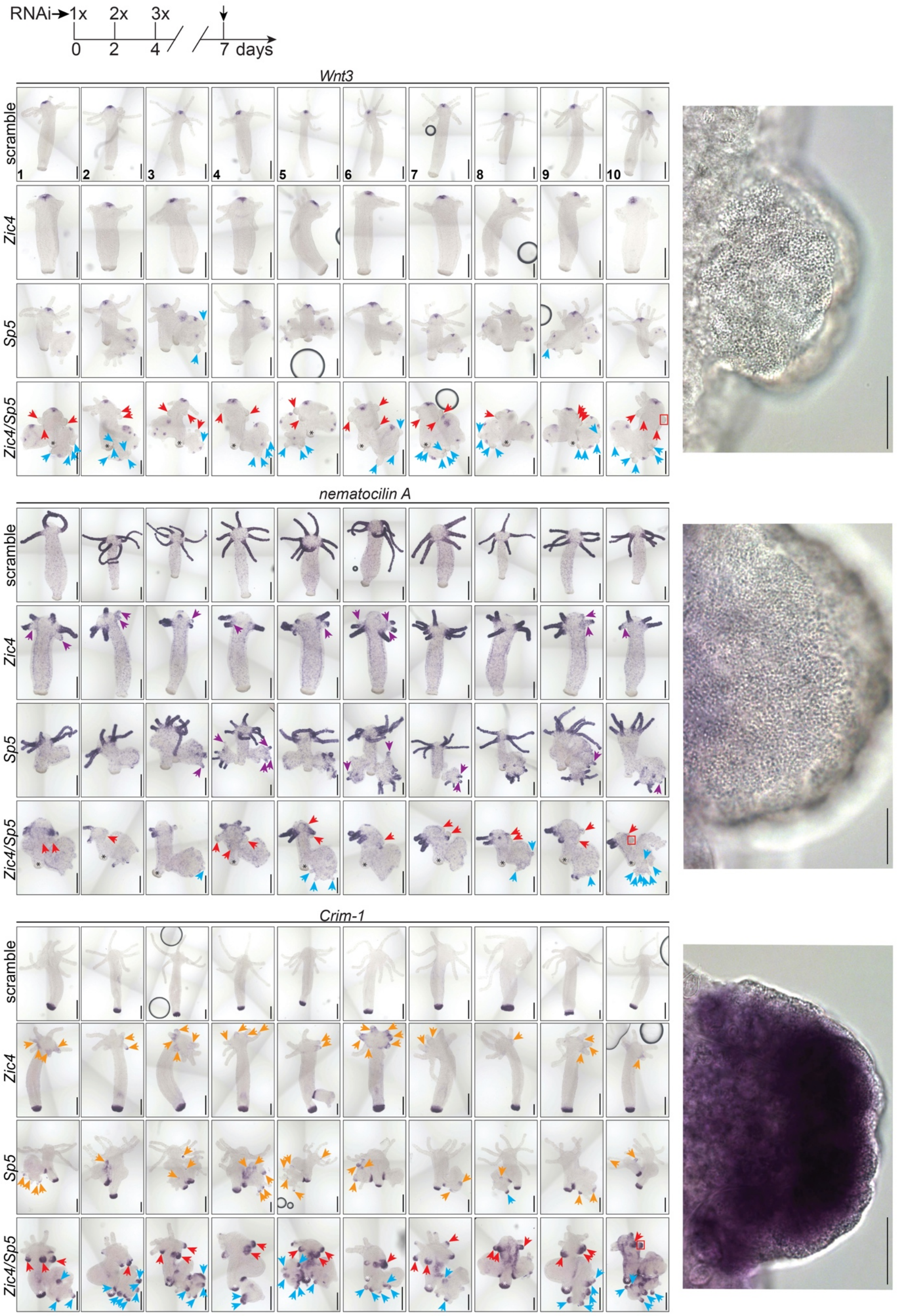


**Figure Supplement 20. Ectopic basal disc formation upon *Zic4*, *Sp5* or *Zic4*/*Sp5*(RNAi) in *Hv_Basel***

*Wnt3, nematocilin A* and *Crim-1* expression in intact *Hv_Basel* animals electroporated three times with a scramble siRNA or *Zic4, Sp5*, *Zic4/Sp5* siRNAs and fixed three days after RNAi3. The silencing of *Zic4* and *Sp5* causes a loss in the expression of the tentacle marker *nematocilin A* (purple arrows) and the up-regulation of the basal disc specific marker *Crim-1* (orange arrows). This effect is enhanced when both genes are silenced together, which results in ectopic basal disc formation. Red arrows point to ectopic basal discs in pre-existing heads and blue arrows to basal discs formed in ectopic heads. Red squares on animal 10 of the *Zic4/Sp5*(RNAi) series indicate enlarged regions on the right.

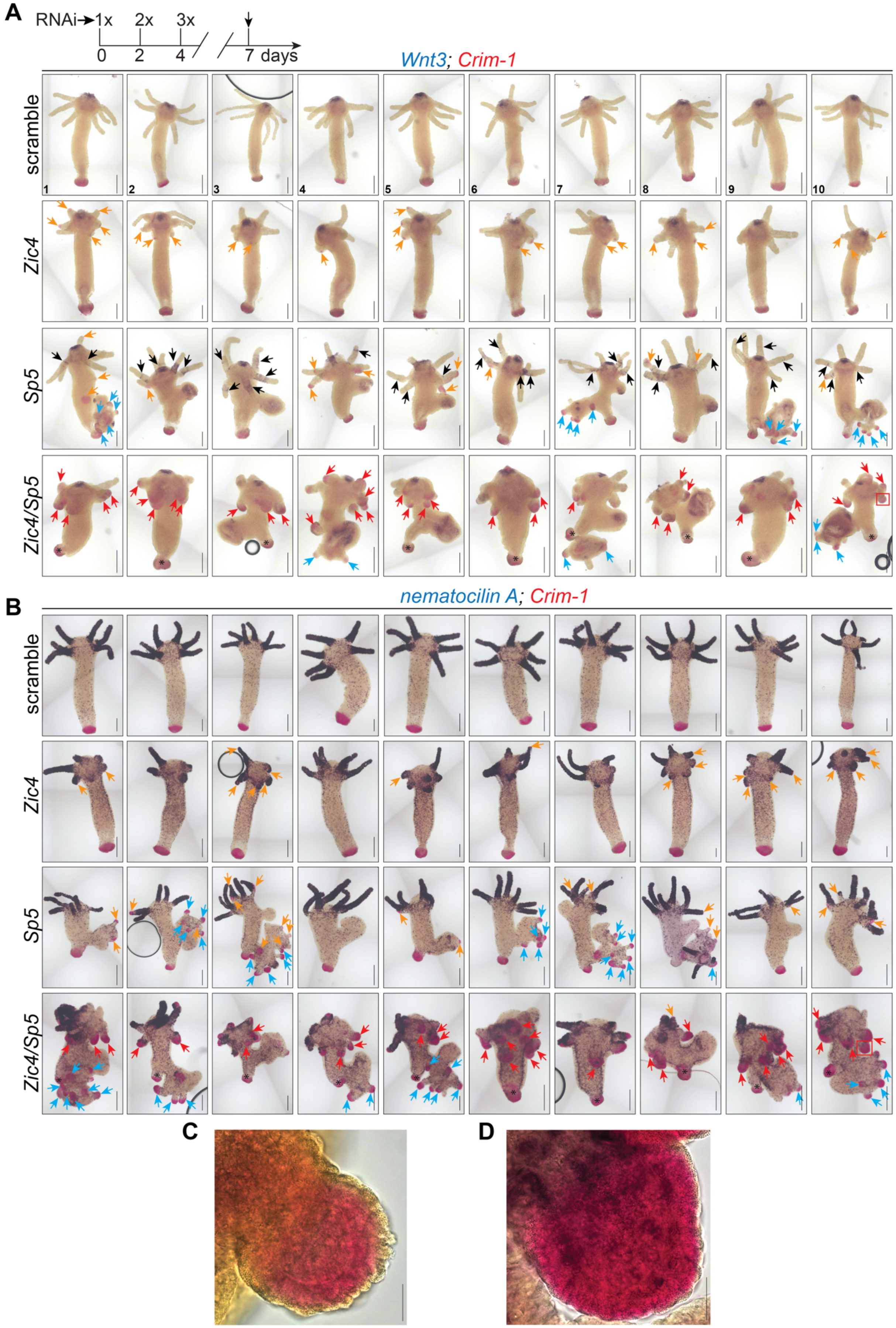
A
RNAi
1x
2x
3x
0
2
4
7 days
Wnt3; Crim-1
scramble
Zic4
Sp5
Zic4/Sp5
1
2
3
4
5
6
7
8
9
10
B
nematocilin A; Crim-1
scramble
Zic4
Sp5
Zic4/Sp5
C
D

**Figure Supplement 21. Ectopic basal disc formation upon *Zic4*, *Sp5* or *Zic4/Sp5*(RNAi) in *Hv_AEP2***

Intact *Hv_AEP2* animals electroporated three times with a scramble siRNA or *Zic4, Sp5*, *Zic4/Sp5* siRNAs were fixed three days after RNAi3 to be co-detected for *Wnt3* (purple) and *Crim-1* (pink) **(A)** or *nematocilin A* (purple) and *Crim-1* (pink) **(B)**. Note the loss of expression of the tentacle marker *nematocilin A* (purple arrows) and the ectopic expression of the basal disc marker *Crim-1* (orange arrows) in tentacles of animals knocked-down for *Zic4*, *Sp5* or *Zic4/Sp5*, effects that are enhanced in the latter condition. Red arrows point to basal tissue differentiating in heads, blue arrows to basal discs differentiating within ectopic structures formed along the body axis in *Sp5*(RNAi) and *Zic4/Sp5*(RNAi) animals; asterisks indicate pre-existing basal discs. Note that *Zic4/Sp5*(RNAi) does not affect the expression of *Wnt3* in the original apical region. For each condition 10 representative animals of an experiment performed in duplicate (n = 10 per replicate) are shown. **(C, D)** Enlarged tentacle of a *Zic4/Sp5*(RNAi) animal (animal 10, red squares) detected for *Wnt3* and *Crim-1* (C) or *nematocilin A* and *Crim-1* (D). Scale bars, 250 µm (A-B); 25 µm (C-D).

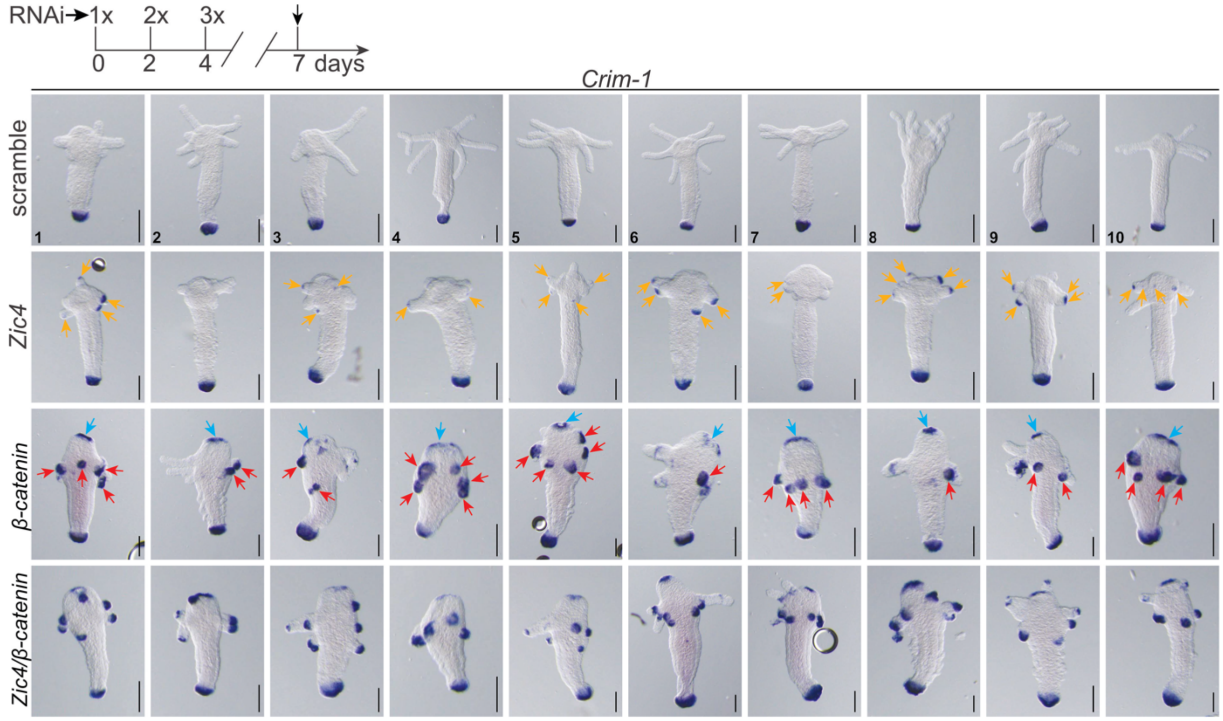


**Figure Supplement 22. Ectopic basal disc formation after *β-catenin*(RNAi) *in Hv_AEP2 animals***

*Crim-1* expression in animals electroporated three times with a scramble siRNA or *Zic4*, *β-catenin* siRNAs and fixed three days later. Note the formation of ectopic basal tissue after knocking-down *Zic4* (orange arrows) or *β-catenin* (red arrows) including at the apical tip (blue arrows). Animals depicted here are representative animals of an experiment performed in duplicate (n = 10 per replicate). Scale bars: 200 µm.

#### (A) EXPRESSION in ECTOPIC STRUCTURES

| | *Hv_Basel* | | | | | | | | | | | |
|---|---|---|---|---|---|---|---|---|---|---|---|---|
| markers | scramble | | | *Zic4*(RNAi) | | | *Sp5*(RNAi) | | | *Zic4/Sp5*(RNAi) | | |
| ***Wnt3*** | 0/53 | 0% | - | 0/57 | 0% | - | 40/40 | 100% | +++ | 55/55 | 100% | +++ |
| ***NemA*** | 0/33 | 0% | - | 0/31 | 0% | - | 26/37 | 70.3% | + | 9/34 | 26.5% | ± |
| ***Crim-1*** | 0/65 | 0% | - | 0/57 | 0% | ± | 30/47 | 63.8% | + | 39/51 | 76.5% | +++ |
| | ***Hv_AEP2*** | | | | | | | | | | | |
| markers | scramble | | | *Zic4*(RNAi) | | | *Sp5*(RNAi) | | | *Zic4/Sp5*(RNAi) | | |
| ***Wnt3*** | 0/21 | 0% | - | 0/18 | 0% | - | 18/21 | 85.7% | ++ | 8/20 | 40% | ++ |
| ***NemA*** | 0/18 | 0% | - | 0/19 | 0% | - | 5/19 | 26.3% | + | 3/19 | 15.8% | ± |
| ***Crim-1*** | 0/39 | 0% | - | 0/37 | 0% | - | 18/40 | 45% | ++ | 17/39 | 43.6% | ++ |

#### (B) EXPRESSION in ORIGINAL HEADS

| | ***Hv_Basel*** | | | | | | | | | | | |
|---|---|---|---|---|---|---|---|---|---|---|---|---|
| markers | scramble | | | *Zic4*(RNAi) | | | *Sp5*(RNAi) | | | *Zic4/Sp5*(RNAi) | | |
| ***Wnt3*** | 53/53 | 100% | +++ | 57/57 | 100% | +++ | 40/40 | 100% | +++ | 55/55 | 100% | +++ |
| ***NemA*** | 33/33 | 100% | +++ | 31/31 | 100% | ++ | 37/37 | 100% | +++ | 34/34 | 100% | + |
| ***Crim-1*** | 0/65 | 100% | - | 48/57 | 84.2% | ± | 5/47 | 10.6% | ± | 51/51 | 100% | +++ |
| | ***Hv_AEP2*** | | | | | | | | | | | |
| markers | scramble | | | *Zic4*(RNAi) | | | *Sp5*(RNAi) | | | *Zic4/Sp5*(RNAi) | | |
| ***Wnt3*** | 21/21 | 100% | +++ | 18/18 | 100% | +++ | 21/21 | 100% | +++ | 20/20 | 100% | +++ |
| ***NemA*** | 18/18 | 100% | +++ | 19/19 | 100% | ++ | 19/19 | 100% | +++ | 19/19 | 100% | + |
| ***Crim-1*** | 0/39 | 100% | - | 32/37 | 86.5% | ± | 20/40 | 50% | ± | 39/39 | 100% | +++ |
| | ***Hv_AEP2*** | | | | | | | | | | | |
| markers | scramble | | | *Zic4*(RNAi) | | | β-*catenin*(RNAi) | | | *Zic4*/β-*catenin* (RNAi) | | |
| ***Crim-1*** | 0/13 | 0% | - | 11/17 | 64.7% | + | 17/17 | 100% | +++ | 17/17 | 100% | +++ |

**Figure Supplement 23. *Wnt3, NemA* and *Crim-1* expression in ectopic apical structures (A) or in pre-existing heads (B) when animals are knocked-down for *Zic4, Sp5, Zic4/Sp5, β-catenin* or *Zic4/β-catenin***

This table shows the number of *Hv_AEP2* or *Hv_Basel* animals knocked-down for *Zic4, Sp5, Zic4/Sp5, β-catenin* or *Zic4/β-catenin* expressing the hypostome marker *Wnt3*, the tentacle marker *NemA*, or the basal disc marker *Crim-1* either in the ectopic structures they form 3 days post-EP3 (A), or in the original head 3 days post-EP3 (B). EP3: third electroporation of siRNAs. The intensity of transcript staining was characterized as undetected (-); weak (±), low (+), intermediate (++) or strong (+++).

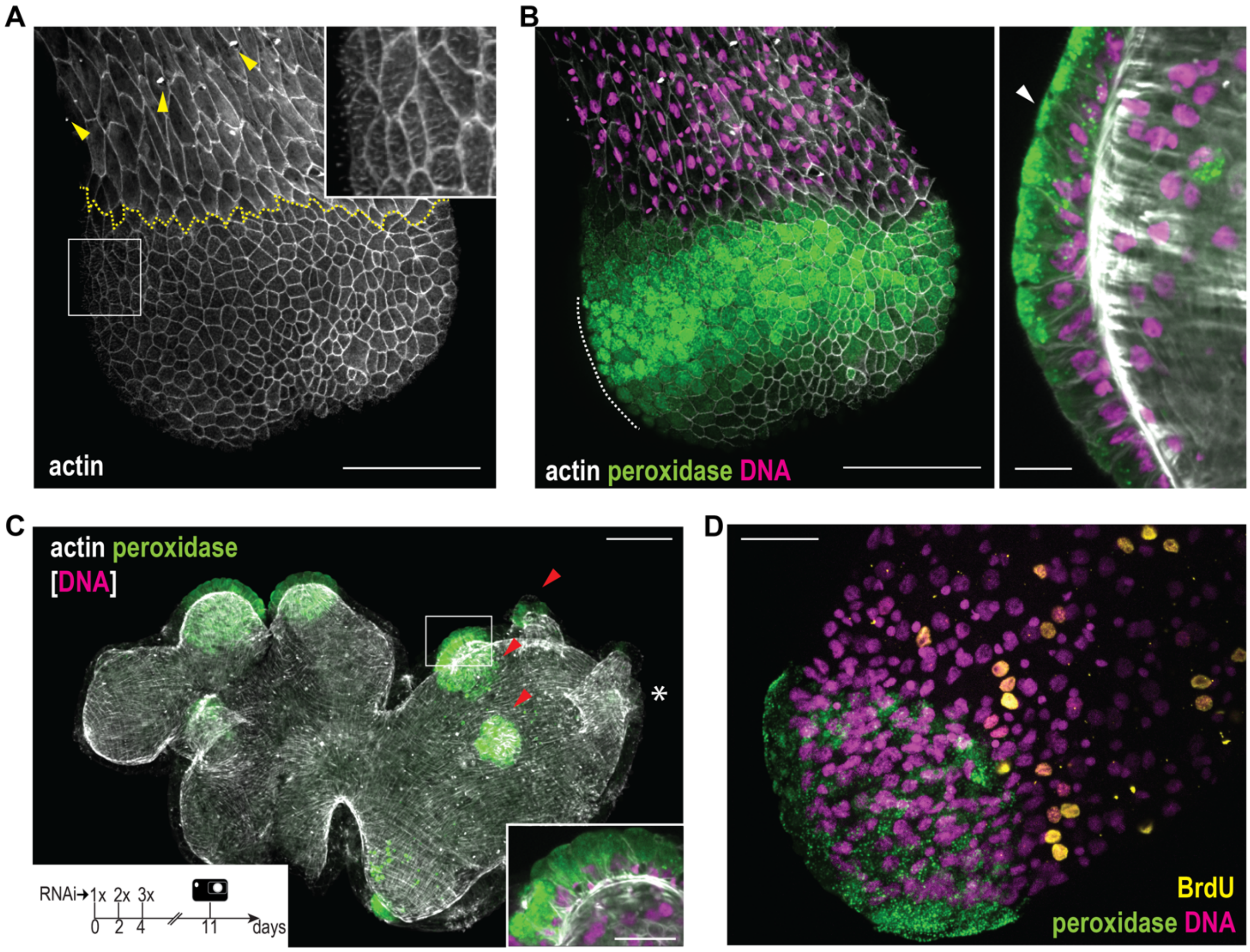


**Figure Supplement 24. Anatomy of basal disc and fully transformed tentacles of *Hv_AEP2***
(**A**) The distinctive morphology of basal disc cells as shown by phalloidin staining (standard deviation Z-projection). Note the typical ciliated cells (inset) and the basal disc area being completely free of nematocytes (yellow arrowheads). Yellow dotted line marks the basal disc boundary. (**B**) High levels of peroxidase activity are typical for basal disc cells. The panel shows a Z-projection of tissue surface layers, while the enlargement shows a midplane optical section. Note the characteristic inverted triangle shape of the basal disc mucous cells with apically concentrated peroxidase granules (white arrowhead) and basally positioned nucleus. (**C**) A representative *Zic4/Sp5*(RNAi) animal 7 days after the last electroporation. The mouth position in the original head is indicated with an asterisk. The original tentacles (red arrowheads) are now almost completely transformed to foot-like structures. Inset shows the cellular morphology in an optical section of one of the transformed original tentacles. (**D**) In a normal basal disc, the peroxidase mucous cells are BrdU negative. Scale bars: 50 µm (A-D), 20 µm (enlargements in B and C).

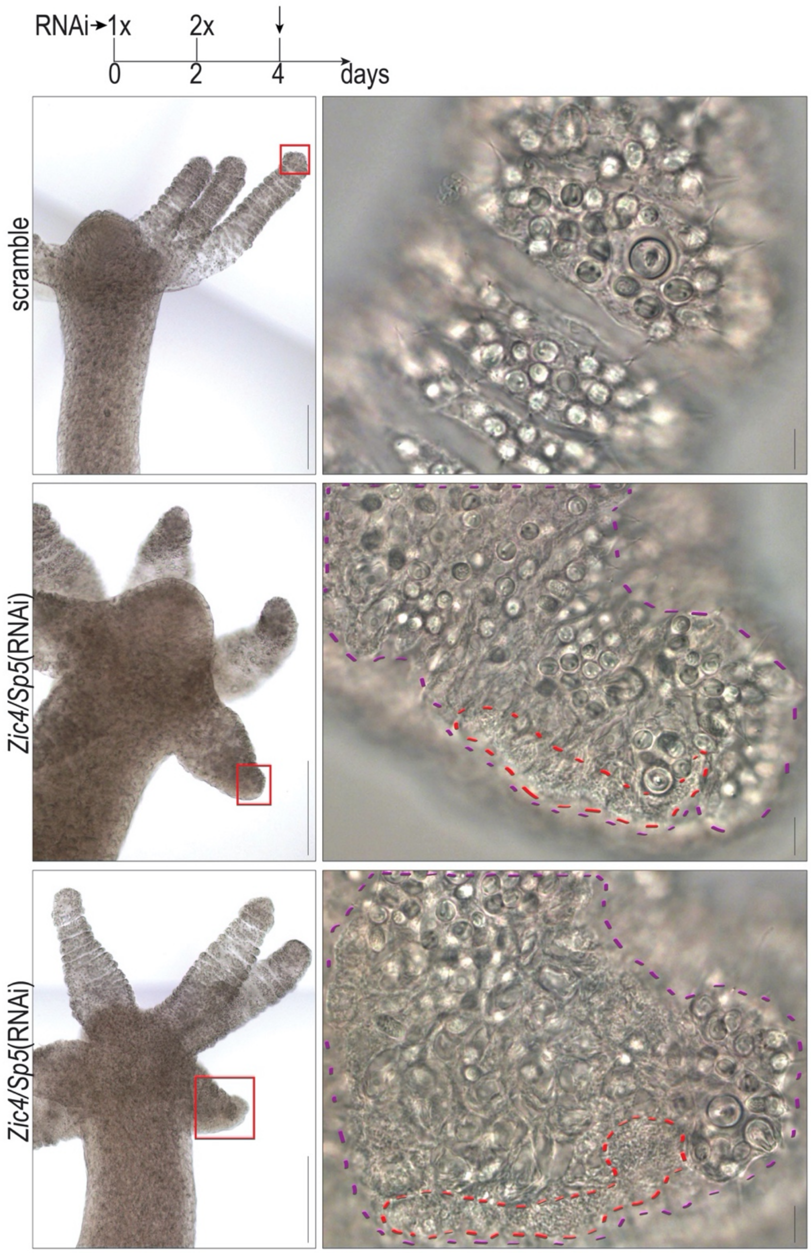


**Figure Supplement 25. Dedifferentiation of tentacle battery cells in *Zic4/Sp5*(RNAi) *Hv_AEP2* animals**

Intact *Hv_AEP2* animals electroporated twice with a scramble siRNA or *Zic4/Sp5* siRNAs were fixed and imaged 2 days post-EP2. Regions of nematocyte degeneration are encircled in purple and tentacle battery cells containing basal-specific acidic mucopolysaccharide (MPS) droplets in red. Enlarged regions of tentacles are indicated with red squares. Two representative *Zic4/Sp5*(RNAi) animals are shown from an experiment performed in triplicate (n=20 animals for each replicate). Scale bars: 200 µm; 10 µm for enlargements.

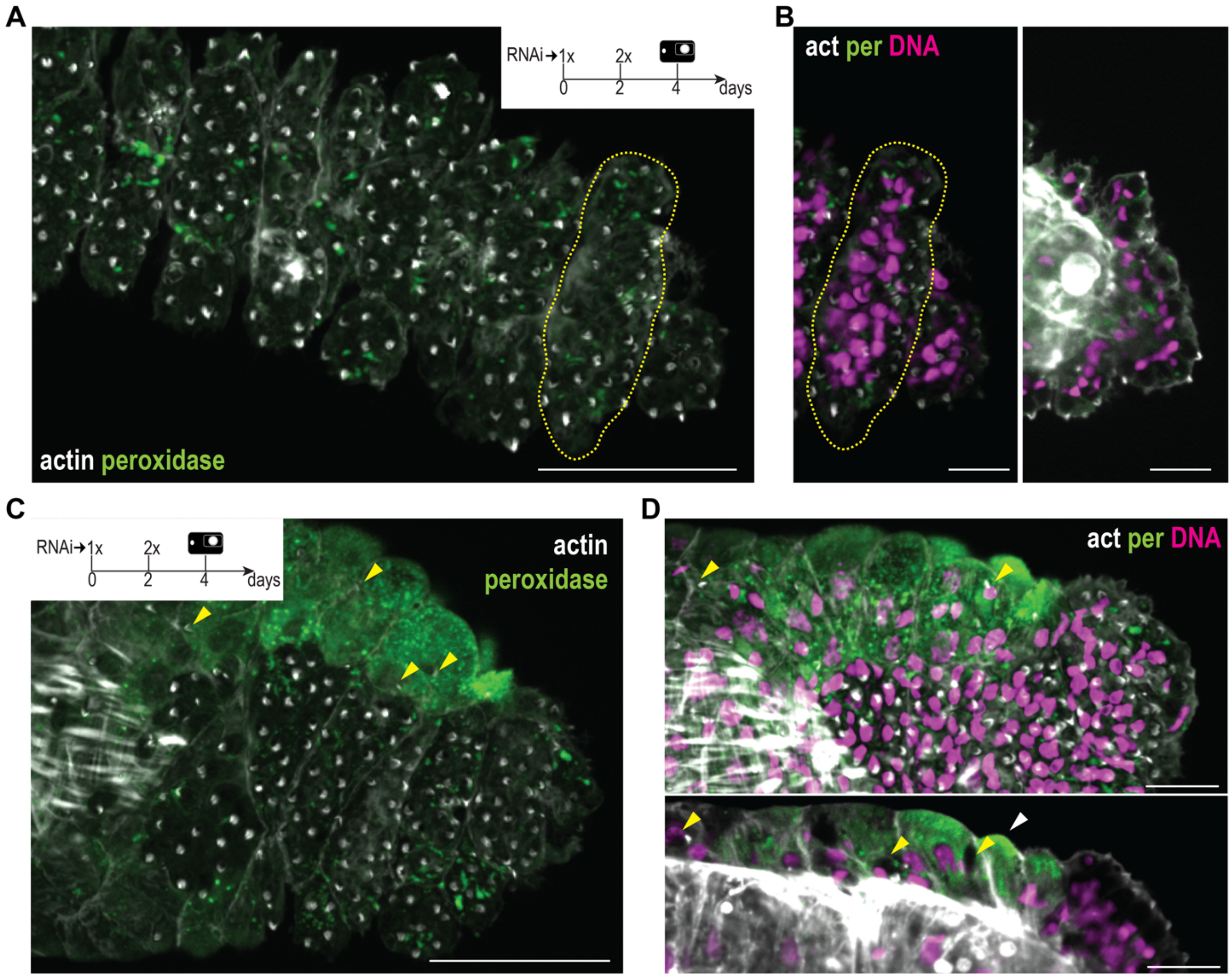


**Figure Supplement 26. Peroxidase activity in TBCs of *Zic4/Sp5*(RNAi) *Hv_AEP2* animals**

(**A**) Phalloidin-stained tentacle after scramble RNAi, negative for peroxidase. Note the oblong battery cells (yellow dotted line) filled with nematocytes, which are visible by their apical actin staining. Standard deviation Z-projection. (**B**) Details of the battery cell structure from (A) including nematocyte nuclei. Z-projection (left) and an optical section through the tentacle middle plane (right). The cell encircled in the yellow dotted line is identical to the one in (A). (**C**) Phalloidin-stained tentacle upon *Zic4/Sp5*(RNAi). The peroxidase positive transformed region harbors only a few nematocytes (yellow arrowheads), compared to the untransformed portion below. Standard deviation Z-projection. (**D**) Details of the boundary between the transformed and intact region from (C), including nuclear staining. Z-projection (upper panel) and an optical section through the tentacle middle plane (lower panel) are shown. Yellow arrowheads indicate retained nematocytes. Scale bars: 50 µm (A-C), 20 µm (B-D).

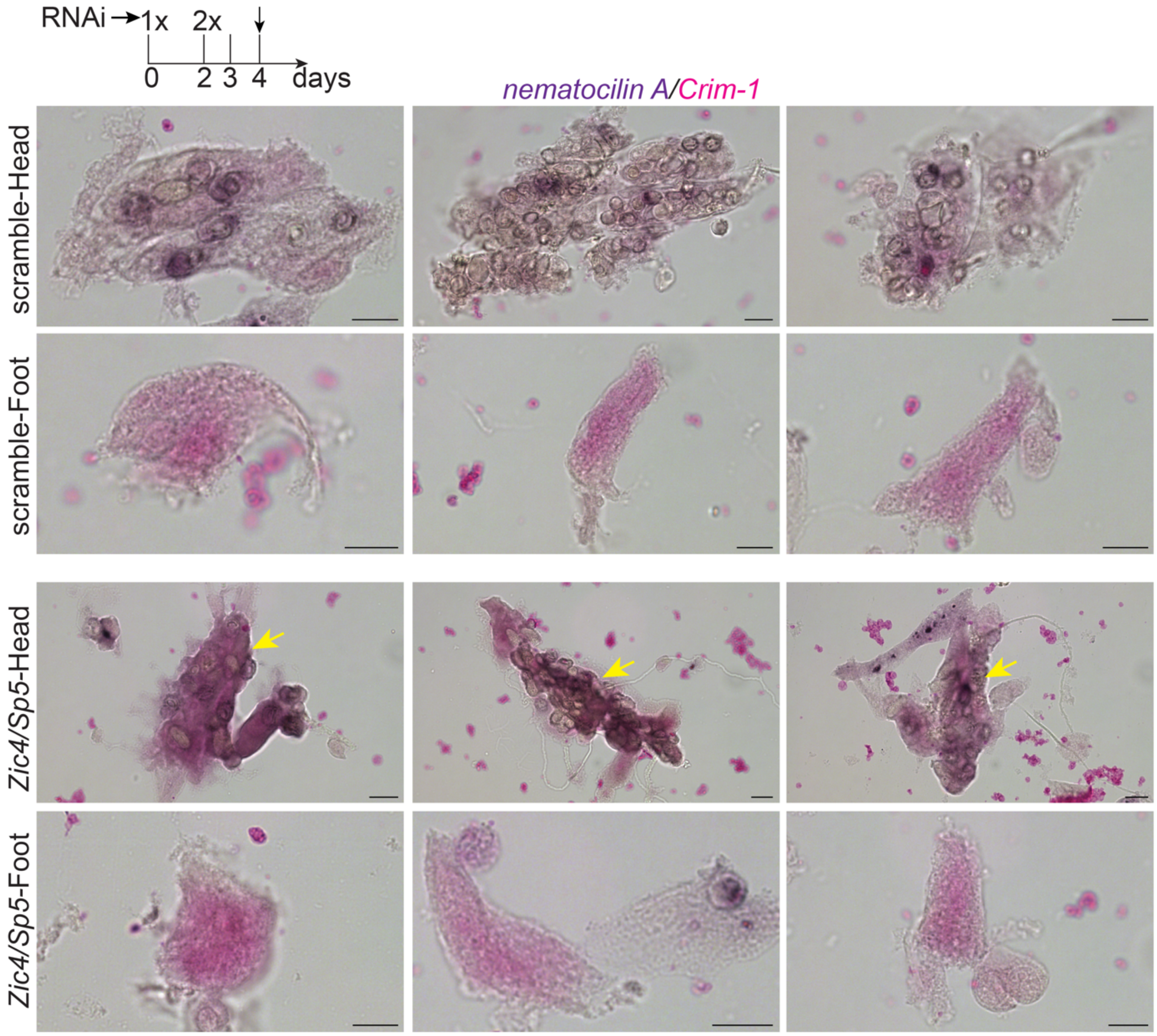


**Figure Supplement 27.** ***Nematocilin A*** **and** ***Crim-1*** **expression detected on macerates**

*Hv_AEP2* animals were electroporated two times with a scramble siRNA or a mix of *Zic4/Sp5* siRNAs and head and foot tissue macerated two days later. Note the co-expression of *NemA* (purple) and *Crim-1* (pink) in tentacle battery cells (yellow arrows) of *Zic/Sp5*(RNAi) animals. Scale bars: 10 µm.

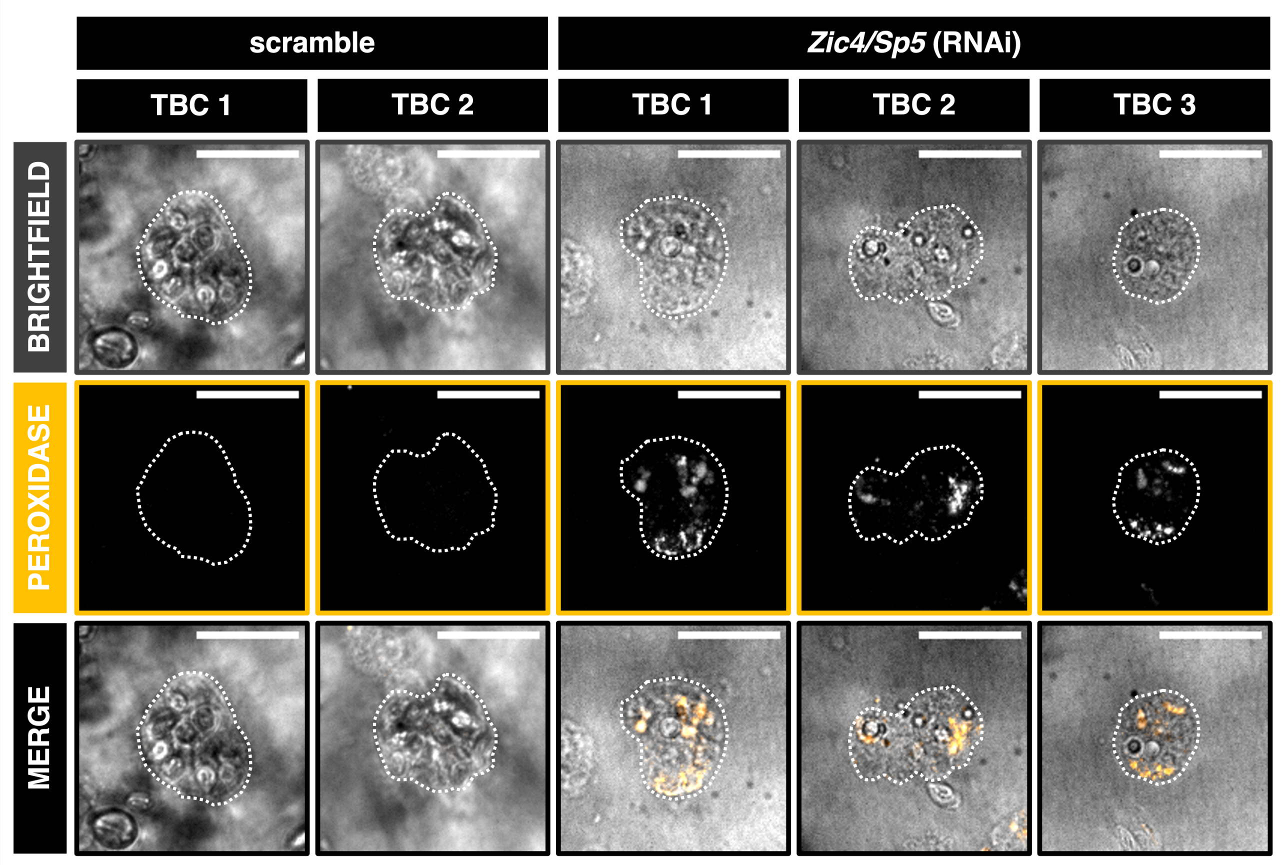


**Figure Supplement 28. Peroxidase activity in isolated Tentacle Battery Cells (TBCs) of *Zic4*/*Sp5*(RNAi) animals**

FUCCI-eGFP *Hydra* were electroporated twice with scramble or a mix of *Zic4/Sp5* siRNAs, then two days post-EP2, the apical regions were dissected, and trypsin macerated. Peroxidase activity in TBCs was subsequently assayed using Tyramide and TBCs were imaged using spinning disk confocal microscopy. Shown are representative TBCs from scramble and *Zic4/Sp5*(RNAi) animals with brightfield channel (single z slice), far red channel corresponding to peroxidase detected with Alexa Fluor™ 647 Tyramide. All images were acquired with the same exposure time and min-max intensities were adjusted similarly 250-487, max projection, and then merged. TBC outlines are marked with a white dashed line. Scale bars: 25 µm.

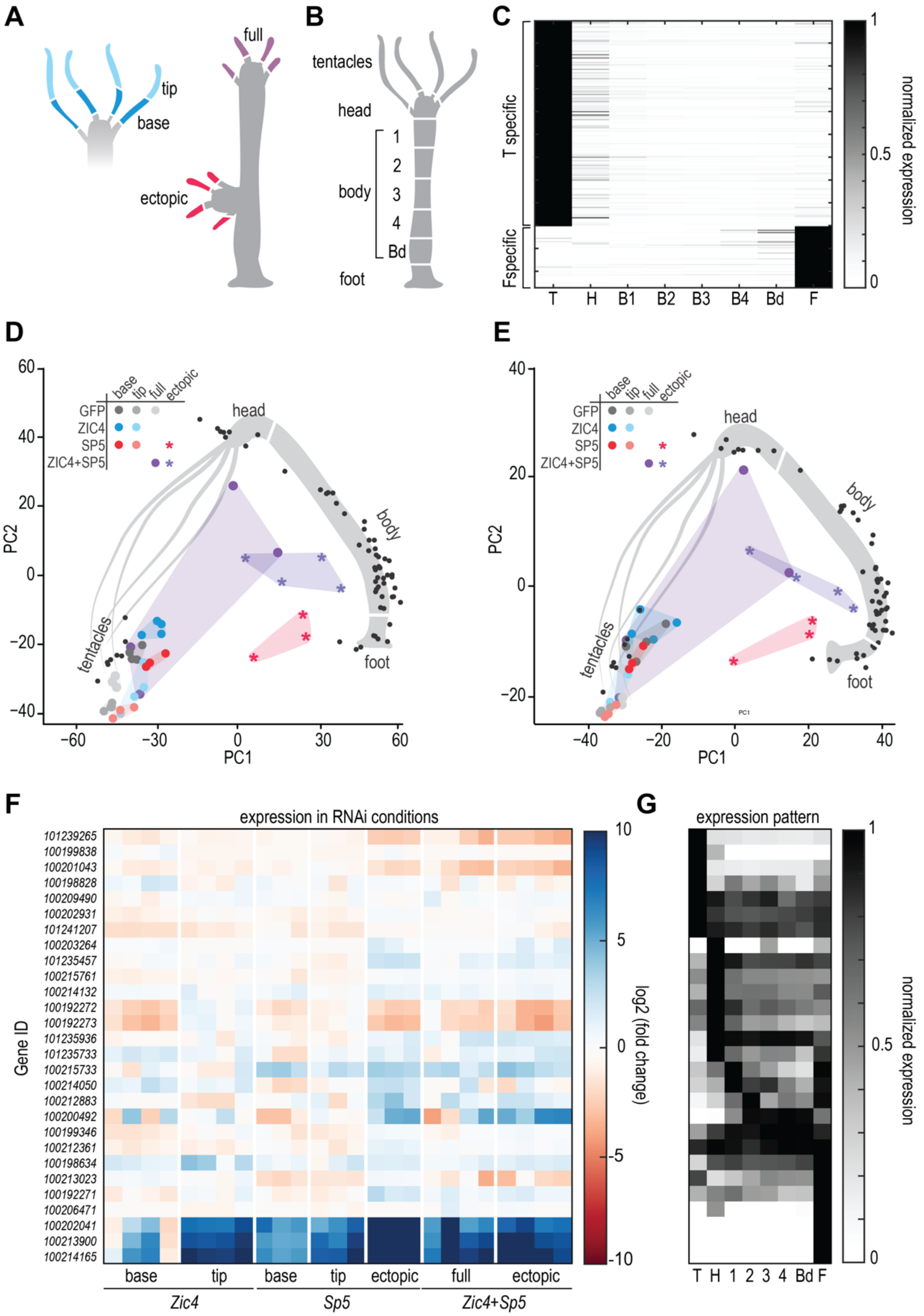


**Figure Supplement 29. RNA sequencing of tentacles from *Zic4*(RNAi) and/or *Sp5*(RNAi) *Hm-105* animals**

(**A**) Scheme showing the way in which the tentacle samples were obtained. When possible, the tentacles of the original head were cut into a proximal (base) and distal (tip) portion. Corresponding pieces were pooled per animal. (**B**) Scheme of the different body parts that were sequenced in the positional RNA-seq (Ferenc et al. 2021). (**C**) Expression pattern of tentacle and basal markers in different body parts. For each gene, the values are normalized to the position with highest expression. T specific: tentacle specific; F specific: basal disc specific. (**D, E**) Projection of the RNAi tentacle samples into the PCA space created from the positional dataset, either using all genes variable among the body parts (D), or using only genes expressed in the interstitial cells and variable among the body parts (E). (**F**) Heatmap of the expression of all *Hydra* peroxidase genes across the RNAi conditions, showing the specific upregulation of a small subset rather than a general stress response. (**G**) Expression profile of the *Hydra* peroxidase genes in (F) along the body, taken from the positional RNA-seq datasets, which indicates that the upregulated peroxidases are basal-specific. T: tentacle; H: heads; Bd: budding zone; F: foot; regions 1, 2, 3, 4 as described in panel B.

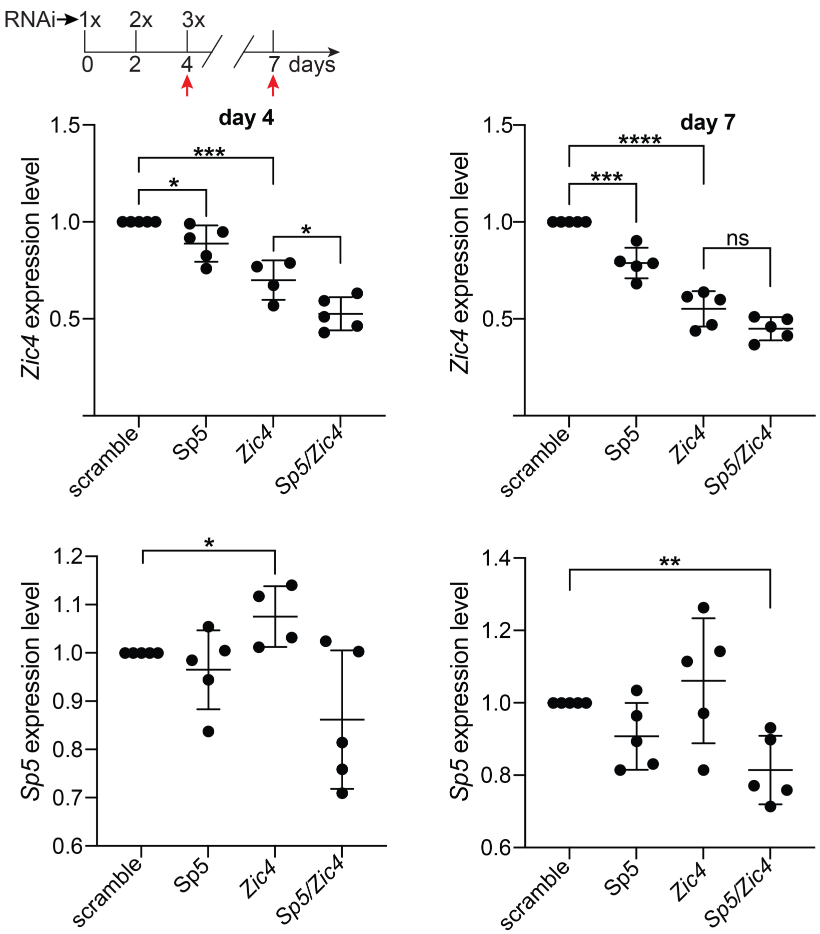


**Figure Supplement 30. Q-PCR analysis of *Zic4* and *Sp5* expression in *Zic4*(RNAi), *Sp5*(RNAi) and *Zic4*/*Sp5*(RNAi) *Hv_AEP2* animals**

Animals were electroporated with a scramble siRNA or siRNAs targeting *Zic4* and *Sp5* and RNA extracted from head tissue on day 4 and day 7, respectively. *≤ 0.05; **≤ 0.01; ***≤0.001; ****≤ 0.0001. Error bars indicate SDs.

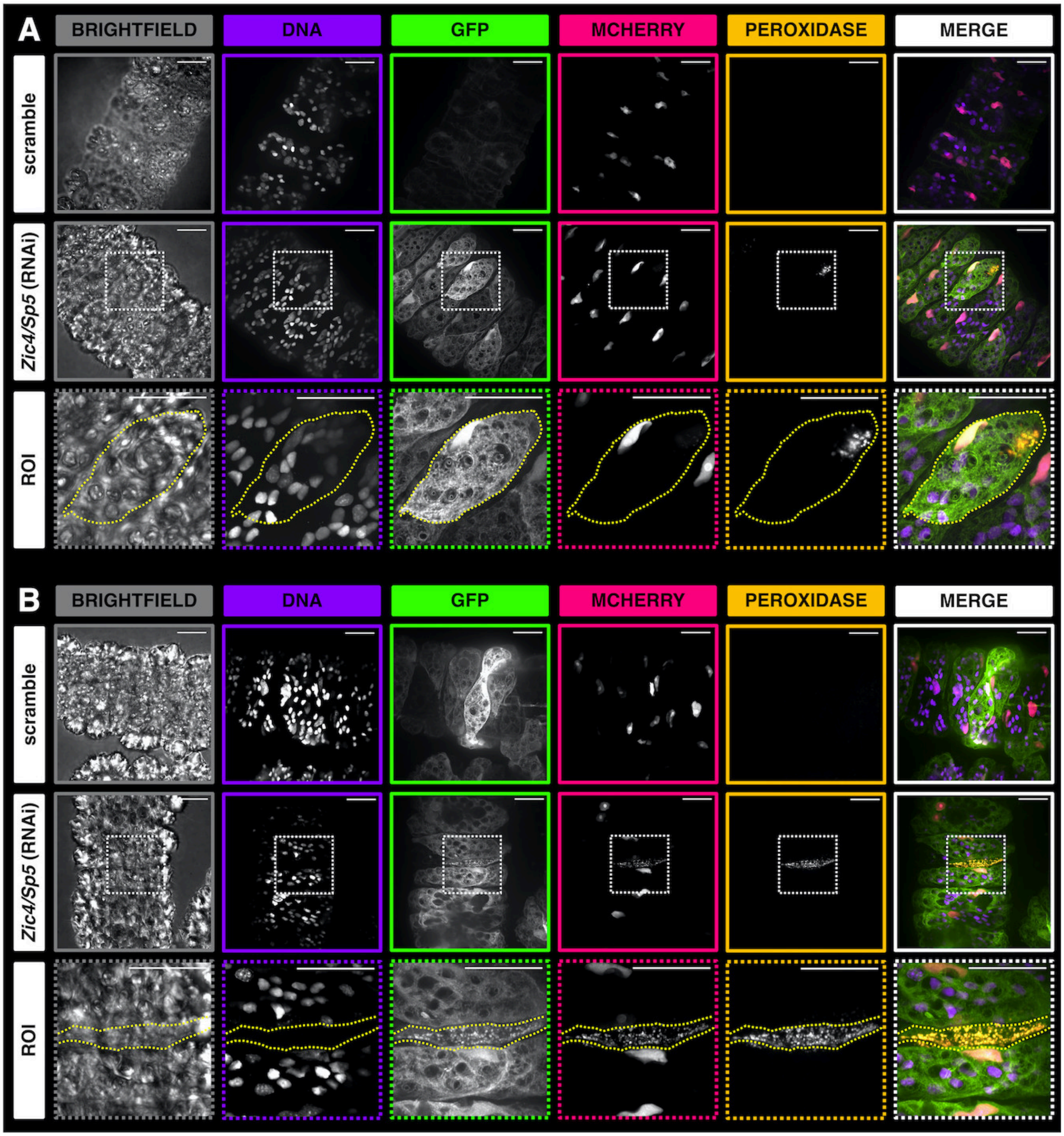


**Figure Supplement 31. Tentacle Battery Cells (TBCs) showing early (A) or advanced (B) signs of transdifferentiation in *Zic4/Sp5* (RNAi) animals**

Intact FUCCI-eGFP *Hydra* expressing Geminin-GFP and CTD-mCherry were electroporated twice with scramble or *Zic4/Sp5* siRNAs, then fixed and treated with Tyramide for detection of peroxidase activity two days post-EP2. Shown are representative tentacles from scramble and *Zic4/Sp5*(RNAi) animals detected with brightfield channel, DNA channel for DAPI-stained nuclei, GFP channel for fixed Geminin-GFP fluorescence marking the cell body and epithelial nuclei, mCherry channel for fixed CTD-mCherry fluorescence marking epithelial nuclei, far red channel for peroxidase activity detected with Alexa Fluor™ 647 Tyramide, and their merge. In each sample, a region-of-interest (ROI) around a TBC that shows early (A) or advanced (B) signs of transdifferentiation, is highlighted with TBC boundaries marked with a yellow dashed line. Samples were imaged via spinning disk confocal microscopy with their respective controls on the same day using the same settings, except for the red channel of (A) where the exposure times were 900 ms and 100 ms for the scramble and *Zic4/Sp5*(RNAi) conditions respectively. Z-stacks were max projected and min-max fluorescence intensities were adjusted similarly between *Zic4/Sp5*(RNAi) and its respective control, except for the red channel of (A). Note that at late-stage of transdifferentiation (B), when the peroxidase activity is higher after *Zic4/Sp5*(RNAi), the Tyramide signal bleeds through the other channels. Scale bars: 25 μm.

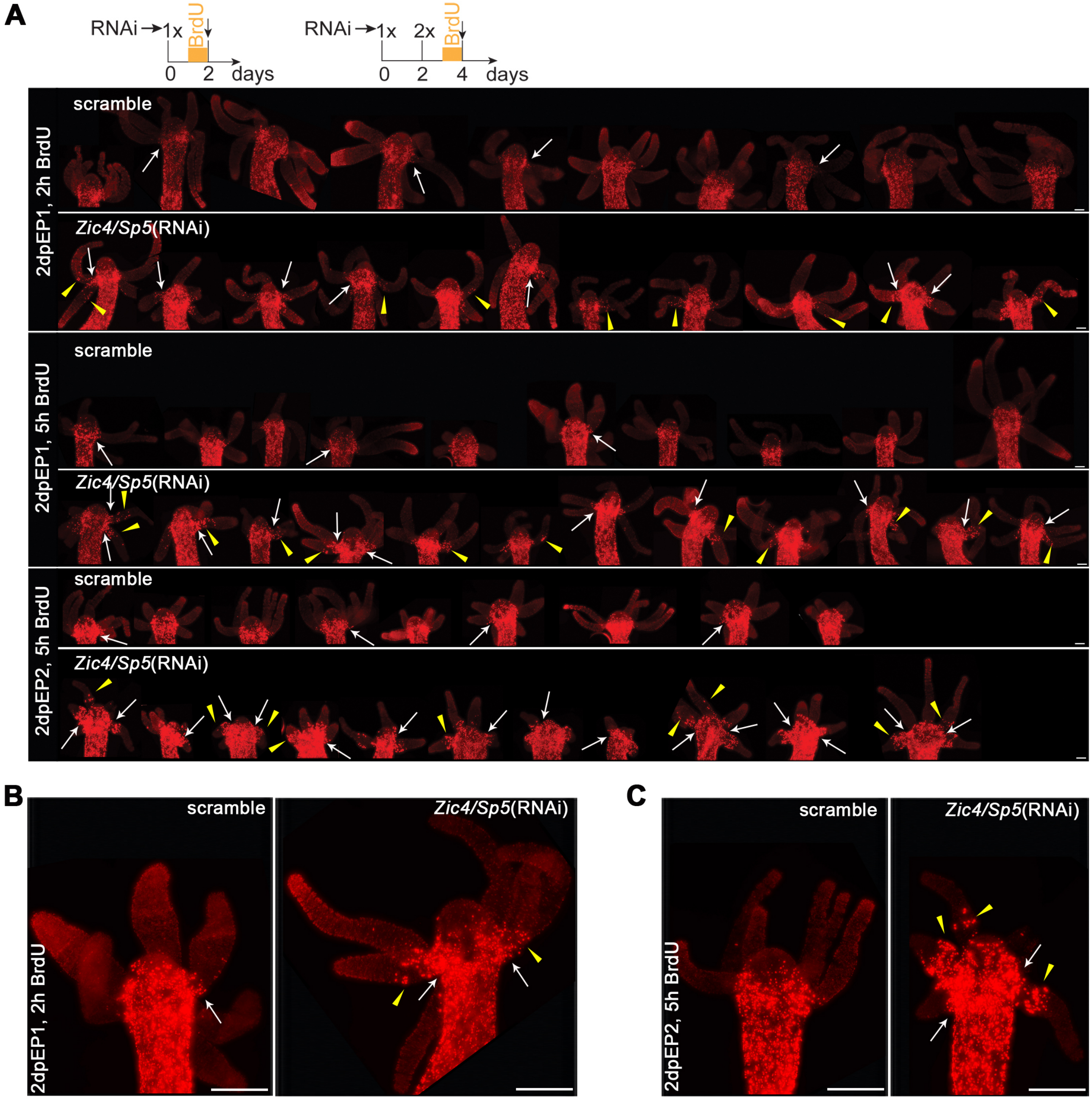


**Figure Supplement 32. Cell proliferation in the apical region after *Zic4/Sp5*(RNAi)**

**(A)** FUCCI-eGFP transgenic *Hydra* electroporated twice with scramble or *Zic4/Sp5* siRNAs were incubated for 2h or 5h with BrdU two days after EP1 (2dpEP1) or after EP2 (2dpEP2) and immediately immunostained for BrdU (red) and GFP. Only the red fluorescence is shown. Each image represents a maximum projection of a 30-40 µm Z-stack. The animals were pictured at 250 msec exposure time except the *Zic4/Sp5*(RNAi) 2dpEP1 and 2dpEP2 conditions that were pictured at 150 msec exposure. To highlight the BrdU+ cells in the tentacles, all images were adjusted in the same way in Photoshop. White arrows and yellow arrowheads point to BrdU+ cells at the root of tentacles and within the tentacles, respectively. Note the increase in proliferating cells in these two regions after *Zic4/Sp5*(RNAi). **(B, C)** Enlarged animals from panel A one day post-EP2 (scramble:

animal 6, *Zic4/Sp5*(RNAi): animal 4) and two days post-EP2 (scramble: animal 3, *Zic4/Sp5*(RNAi): animal 1). Scale bars: 250 µm.

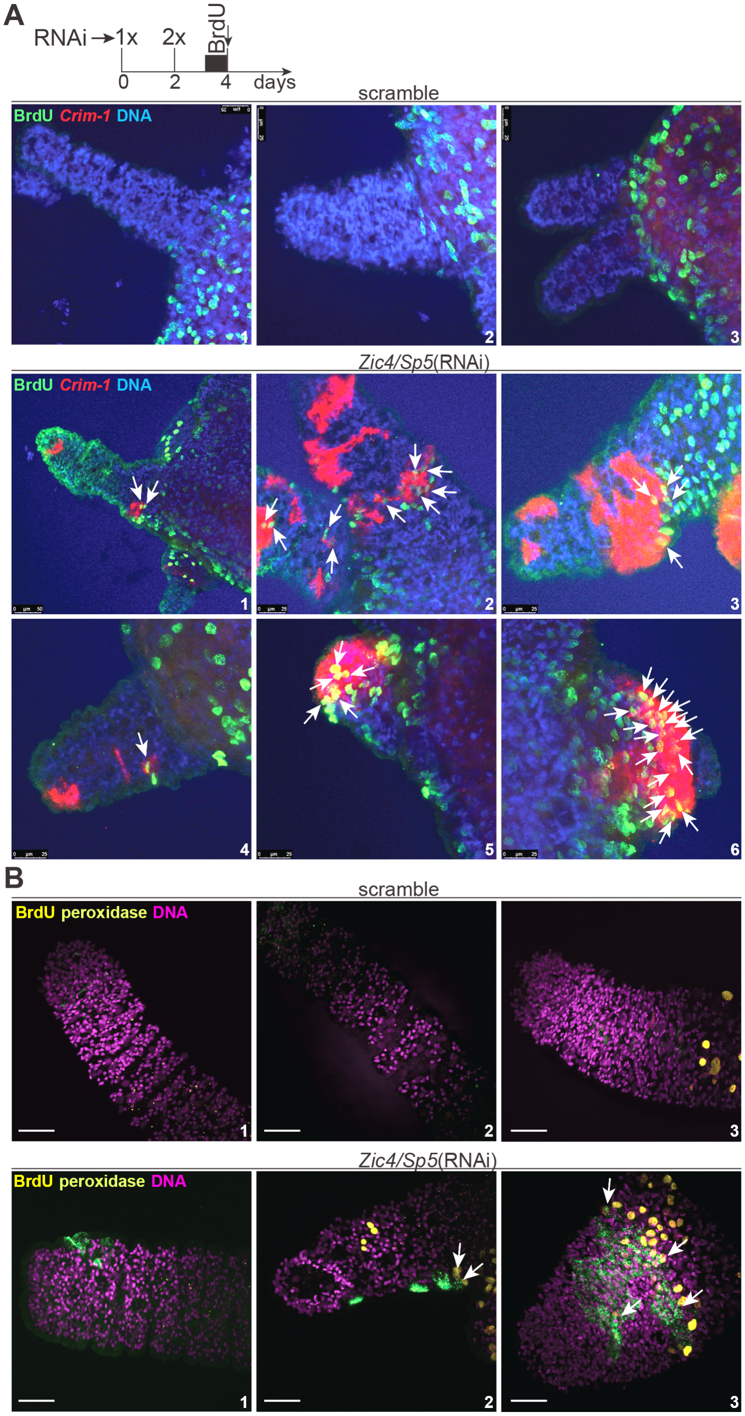


**Figure Supplement 33.** ***Crim-1*** **expression and peroxidase activity in BrdU+ tentacle cells after** ***Zic4/Sp5*****(RNAi)**

Intact *Hv_AEP2* animals were electroporated two times with a scramble siRNA or a mix of *Zic4/Sp5* siRNAs, treated with BrdU for 5 h and fixed on day 4. The expression of *Crim-1* (A) was detected by in *situ* hybridization and peroxidase activity (B) was detected with Tyramide (see

Materials and Methods). White arrows points towards *Crim-1*/BrdU+ cells and peroxidase/BrdU+ cells, respectively. Scale bars: 50 µm for (B).

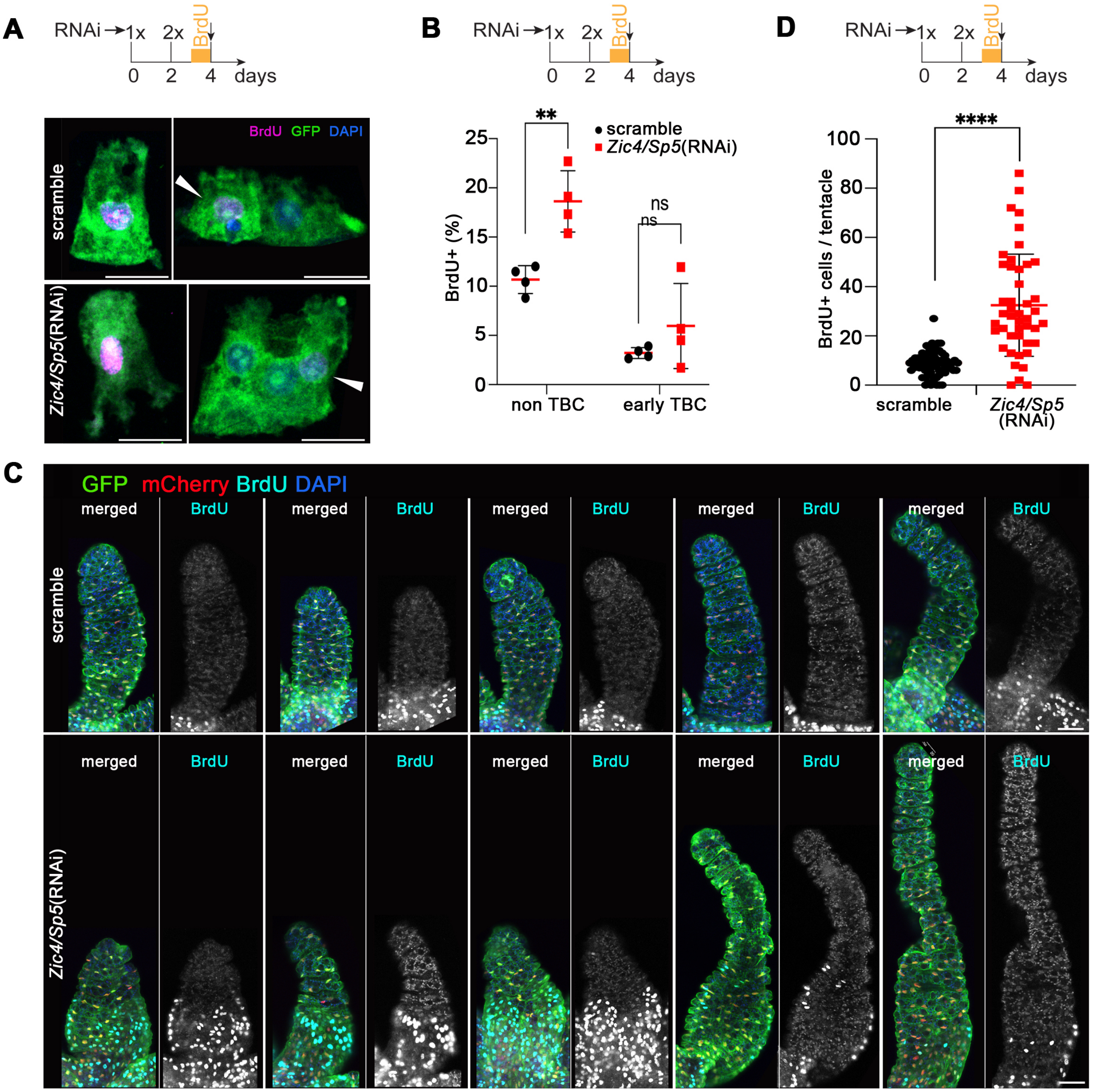


**Figure Supplement 34. Cell cycle re-entry of epidermal epithelial cells after *Zic4/Sp5*(RNAi)**
FUCCI eGFP transgenic *Hydra* that constitutively express the FUCCI eGFP-mCherry construct in epidermal epithelial cells were electroporated twice with scramble or *Zic4/Sp5* siRNAs and exposed two days post-EP2 to BrdU for 5 hours. (**A, B**) The apical region of BrdU-labeled FUCCI animals was cut off immediately after BrdU labeling, macerated and immunostained for BrdU (magenta) and GFP (green). **(A)** Apical BrdU+/GFP+ cells from scramble and *Zic4/Sp5*(RNAi) animals (white arrowheads). The pictures were acquired in the same conditions and the min-max fluorescence intensity of BrdU were adjusted similarly for all images. Scale bars: 25 µm. **(B)** Quantification of

apical BrdU+/GFP+ cells from apical and *Zic4/Sp5*(RNAi) animals. At least 500 GFP+ cells were randomly pictured on each slide and BrdU+ cells were classified as epidermal epithelial cells that are not TBCs, early TBCs that contain up to 9 nematocytes (26). Mature TBCs (over 10 nematocytes) were not represented as too few were BrdU+. At least 230 cells/replicate for each category were counted and the respective percentages of each category are represented, corresponding to one experiment with four replicates. Statistical p-value: **≤ 0.01. Errors bars indicate SD. **(C, D)** Proliferating epithelial cells in tentacles of intact scramble and *Zic4/Sp5*(RNAi) FUCCI-eGFP animals immunodetected for GFP, mCherry and BrdU immediately after BrdU labeling. As above, BrdU+/GFP+ cells correspond to proliferating epithelial cells located in the vicinity of the tentacle root or along the tentacles. **(D)** Number of BrdU+/GFP+ cells in the vicinity of tentacle roots and along the tentacles. Minimum 12 animals/condition were analyzed and tentacles that were superposing partially or totally with other tentacles were excluded. Each point represents a tentacle. Statistical p-value: ****≤ 0.0001. Error bars indicates SD. Scale bars: 50 µm.

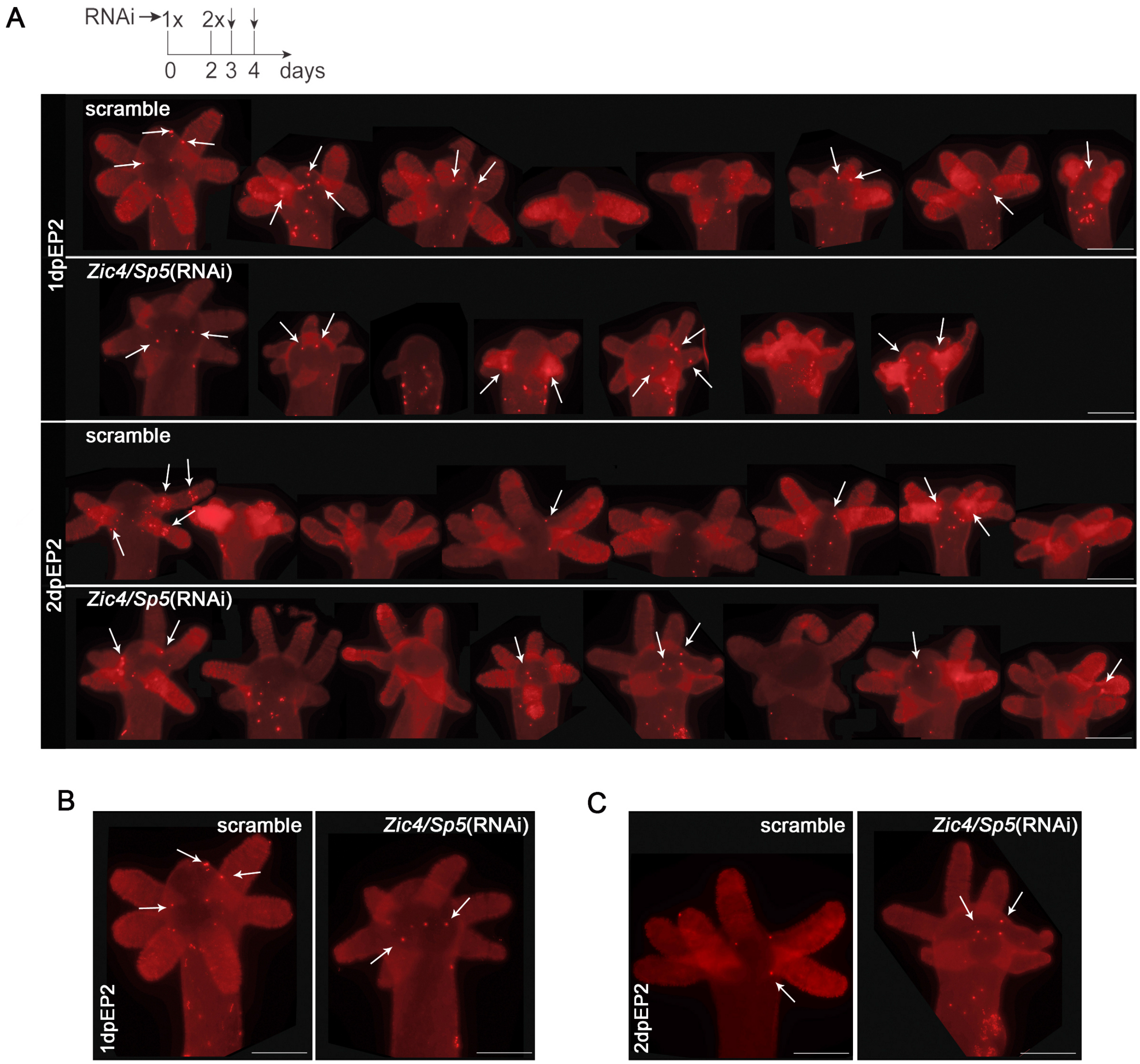


**Figure Supplement 35. Phospho-histone H3 immunostaining of *Zic4/Sp5*(RNAi) animals**

*Hv_AEP2* polyps electroporated with scramble or *Zic4/Sp5* siRNAs were immunostained for phosphohistone H3 (red) one day (1dpEP2) or two days (2dpEP2) post-EP2 **(A)**. Each image represents a maximal projection of a Z-stack covering 30-40 µm. The Z-stacks were acquired at the same exposure time and the intensity of the red fluorescence was increased in the same way in Photoshop. Note the very few cells that enter mitosis in the apical region of scramble and *Zic4/Sp5*(RNAi) animals (white arrows). **(B, C)** Enlarged animals at one day (from left animal 1-scramble, animal 1-*Zic4/Sp5*(RNAi)) and two days (from left animal 4-scramble, animal 5-*Zic4/Sp5*(RNAi)) post-EP2 immunostained for phosphohistone H3. Scale bars: 250 µm.

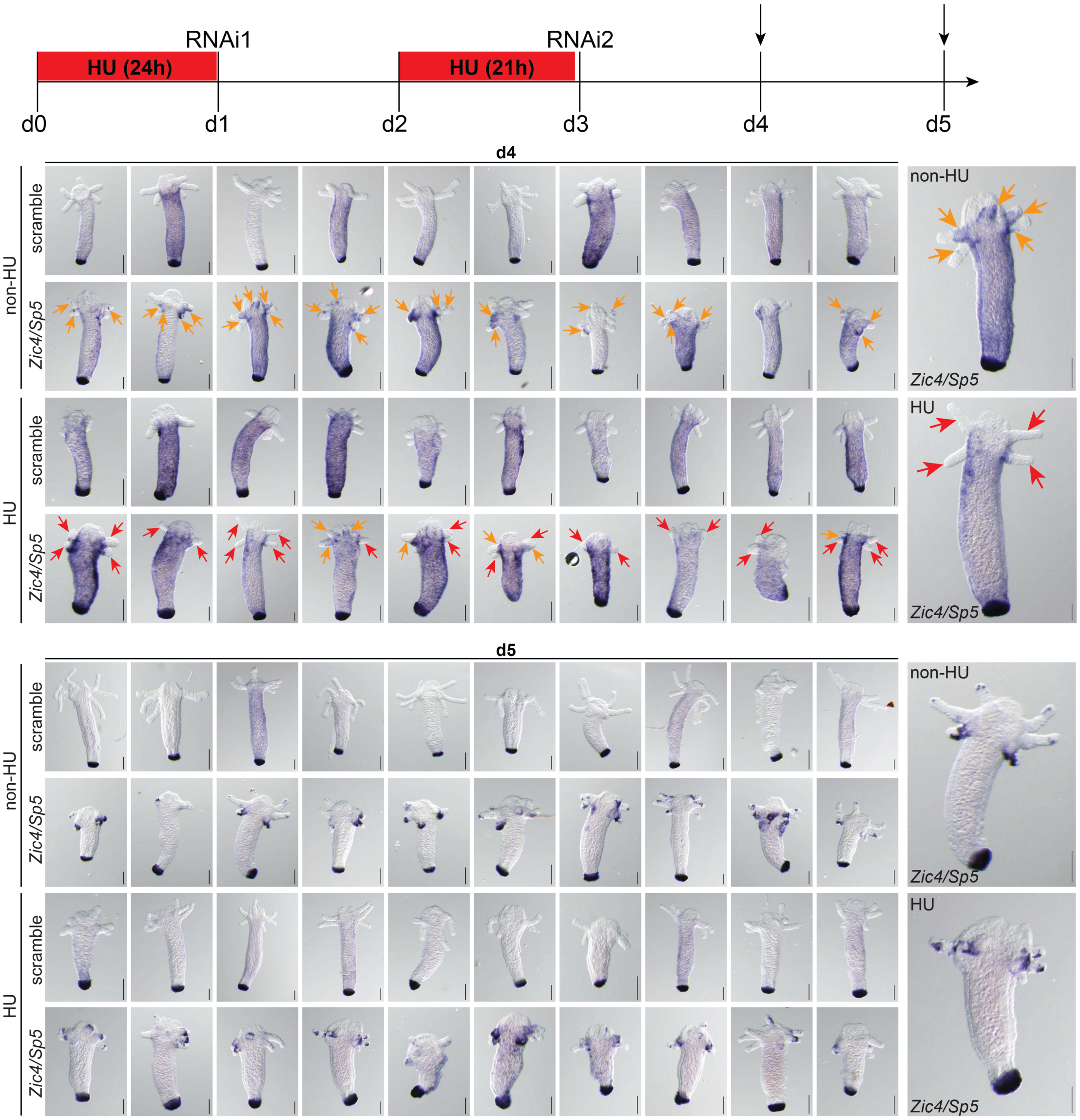


**Figure Supplement 36.** ***Crim-1*** **expression in HU-treated animals**

Intact *Hv_AEP2* animals were electroporated two times with a scramble siRNA or a mix of *Zic4/Sp5* siRNAs and treated with Hydroxyurea (HU) as depicted in the scheme. Animals were processed for *in situ* hybridization on day 4 and 5, respectively to detect the expression of *Crim-1*. Note the reduction of *Crim-1* expression in the tentacles of HU-treated *Zic4/Sp5*(RNAi) animals on day 4 compared to tentacles of non-HU-treated *Zic4/Sp5*(RNAi) animals. Red arrows point towards tentacles lacking *Crim-1* expression and orange arrows point towards *Crim-1* expressing tentacles. Scale bars: 200 µm.

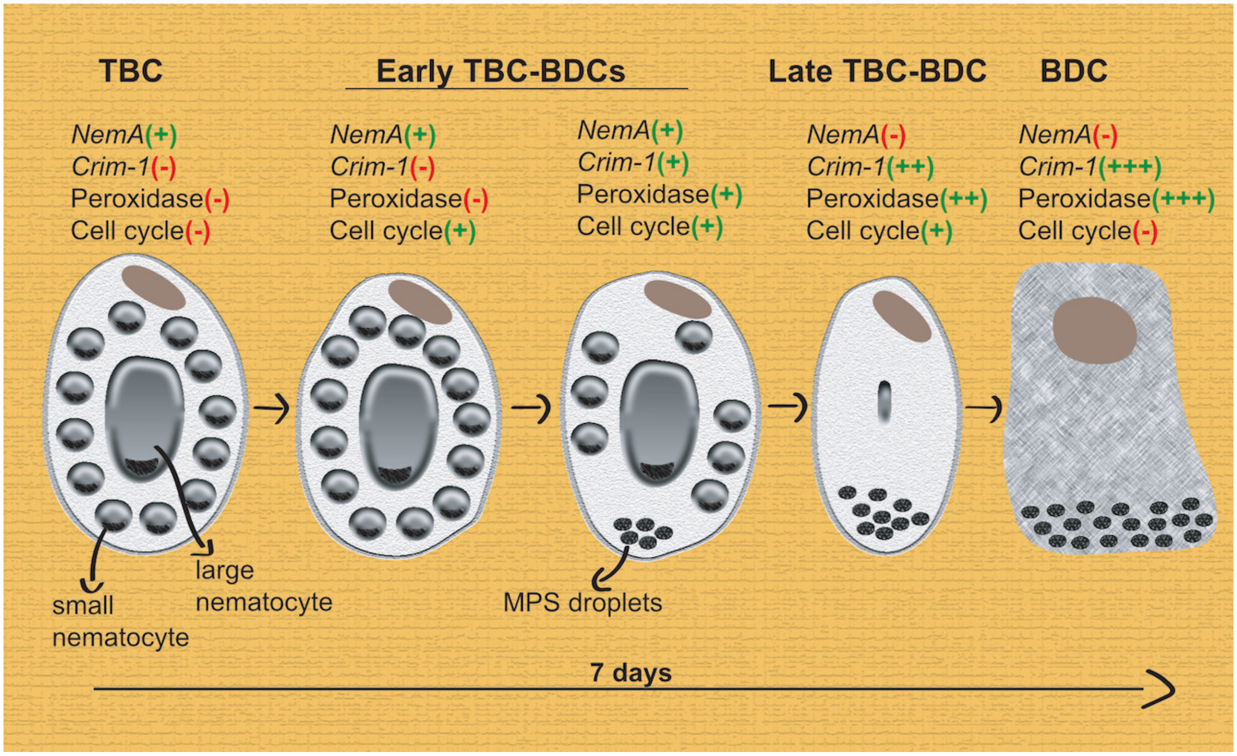


**Figure Supplement 37. Schematic view of the successive stages of transdifferentiation of Tentacle Battery cells (TBC) to Basal Disc cells (BDC).**
